# The Impact of Coronavirus Nsp1 on Host mRNA Degradation Is Independent of Its Role in Translation Inhibition

**DOI:** 10.1101/2024.08.05.606569

**Authors:** Emilie Bäumlin, Dominic Andenmatten, Jonas Luginbühl, Aurélien Lalou, Nino Schwaller, Evangelos D. Karousis

## Abstract

When host cells are infected with coronaviruses, the first viral protein produced is Nsp1. This protein inhibits host protein synthesis and induces host mRNA degradation to enhance viral proliferation. Despite its critical role, the mechanism by which Nsp1 mediates cellular mRNA degradation remains unclear. In this study, we use cell-free translation to address how the host mRNA stability is regulated by Nsp1. We reveal that SARS-CoV-2 Nsp1 binding to the ribosome is enough to trigger mRNA degradation independently of ribosome collisions or active translation. MERS-CoV Nsp1 inhibits translation without triggering degradation, highlighting mechanistic differences between the two Nsp1 counterparts. Nsp1 and viral mRNAs appear to co-evolve, rendering viral mRNAs immune to Nsp1-mediated degradation in SARS-CoV-2, MERS-CoV and Bat-Hp viruses. By providing new insights into the mode of action of Nsp1, our study helps to understand the biology of Nsp1 better and find new strategies for therapeutic targeting against coronaviral infections.

**Significance:**

- Cell-free assays allow the decoupling of Nsp1-mediated translation inhibition from RNA degradation.
- Nsp1 interaction with the ribosome is crucial for mRNA degradation, but active translation is not required.
- SARS-CoV-2 Nsp1 induces mRNA degradation, while MERS-CoV Nsp1 inhibits translation without triggering degradation.
- 5’UTR-specific protection of viral mRNAs from Nsp1 indicates a co-evolutionary adaptation between the two features

## Introduction

Non-structural protein 1 (Nsp1) is the first protein produced during Betacoronavirus infections, playing a key role in hijacking and rendering the host gene expression at the service of the virus. The SARS-CoV-2 Nsp1 protein sequence is divided into an N-terminal domain (NTD) (residues 1-128), a 20-amino-acid-long linker (128-148), and a C-terminal domain (CTD) (148-180) (reviewed in ^1^).

The Nsp1 CTD binds the mRNA entry channel of the 40S ribosomal subunit and leads to host mRNA translation inhibition ^2–5^, allowing, or, in the case of SARS-CoV-2, even promoting the production of viral proteins ^6,7^. This strategy is shared among Betacoronaviruses, such as MERS ^8,9^ and a Bat-infecting coronavirus, Bat-Hp-CoV_Zhejiang2013 referred to as Bat-Hp Coronavirus ^9^. By blocking the mRNA entry channel of host ribosomes, global translation of host mRNAs, including antiviral response mRNAs like IFN, is suppressed ^2,7,10^. Biochemical data support that the Nsp1 NTD also contributes to translation inhibition ^9,11,12^, suggesting that Nsp1 binds the 40S subunit in a bipartite mode of interactions: stably through the CTD with the mRNA entry channel and, more weakly, through the NTD with the decoding centre ^9^.

Apart from inhibiting cellular translation, Nsp1 can also induce host mRNA degradation. The overexpression of Nsp1 in human cells is sufficient to target most of the host mRNAs for degradation ^7,13–16^. Consistent with this observation, SARS-CoV-2-infected cells show an accelerated degradation of mature cytosolic cellular but not viral mRNAs in an Nsp1-dependent manner ^7,17^. Similar to evasion from translation inhibition, the stem-loop 1 (SL1) hairpin correctly positioned at the 5′ end of the viral transcripts is sufficient to protect mRNAs from Nsp1-mediated degradation ^17^. Many observations concerning the mode of action of SARS-CoV-2 Nsp1 are similar to SARS-CoV Nsp1, which shows 84% identity between the two viruses ^18–20^.

Diverse evidence shows that the association of Nsp1 with ribosomes is needed for host mRNA degradation: Nsp1 mutations blocking ribosome binding also inhibit mRNA degradation ^11^, and cell culture experiments show that Nsp1-induced mRNA degradation requires the engagement of 40S or 80S ribosomes on mRNAs ^16^. Moreover, studies in living cells showed that Nsp1 requires the mRNA to interact with ribosomes to induce degradation, in agreement with the observation that upon infection, Nsp1-induced mRNA degradation is more pronounced in actively translated mRNAs ^7^. Furthermore, in Nsp1 transfection experiments, non-translated cytosolic long non-coding RNAs (lncRNAs) resist Nsp1-induced degradation ^16^.

The Nsp1 NTD is crucial for inducing mRNA degradation ^17^. Intriguingly, specific amino acid substitutions at the NTD of SARS-CoV-2 Nsp1 (R125A/K126A) abrogate Nsp1-mediated mRNA cleavage while allowing translation inhibition ^11^. Several lines of evidence suggest that Nsp1 induces mRNA cleavage mainly within the 5΄UTR and the proximal coding region of capped non-viral mRNAs ^17,21^. The host cell exonuclease Xrn1 induces 5΄-3΄ mRNA degradation after SARS-CoV Nsp1 expression ^22^, and interactome studies from SARS-CoV-2 Nsp1 propose that Nsp1 may interact with cellular nucleases ^23^. Based on the hypothesis that Nsp1 recruits cellular nucleases to mediate target mRNA degradation *in vivo* and is directly linked to translation, it has been hypothesized that Nsp1 stimulates ribosome collisions, leading to mRNA degradation relying on ribosome quality control mechanisms (RQC) ^16,24^. In this model, Nsp1 binds to the leading ribosome in a polysome, causing subsequent ribosomes to stall and collide. These collisions are thought to attract host nucleases, which degrade the mRNA. While ribosome collisions are hypothesized to link translation to mRNA degradation, the extent to which Nsp1-induced mRNA degradation relies on ribosome quality control mechanisms (RQC) remains unclear and has not yet been directly tested.

SARS-CoV-2 Nsp1 hinders early host immune responses, lacks homology to human proteins, and has been thoroughly structurally analyzed, making it an ideal drug target. *In silico* studies suggested Montelukast, an FDA-approved compound for asthma treatment and Mitoxantrone, an anticancer drug, which can inhibit the SARS-CoV and SARS-CoV-2 virus entry into cells as potential inhibitors of Nsp1 ^25 26 27^. Furthermore, Ametantrone, structurally similar to Mitoxantrone, was shown to inhibit Nsp1-mediated RNA degradation *in vitro* ^21^.

Exploring whether translation inhibition and mRNA degradation are coupled is crucial for understanding Nsp1 functions. Most approaches to studying the relationship between translation inhibition and host mRNA degradation require monitoring a wide time window after Nsp1 transfection experiments or cellular infections with SARS-CoV-2. For this reason, it has been challenging to determine whether translation inhibition and degradation are coupled activities or if they can occur independently.

Here, we show that in living cells, Nsp1-mediated mRNA decay is not triggered by ribosome collisions and using our human-based cell-free translation system, we show that SARS-CoV-2 Nsp1-induced mRNA degradation can occur independently of translation inhibition. Notably, ribosome binding to Nsp1 is sufficient to trigger degradation without requiring ribosome scanning or translation elongation, and host mRNA degradation is stimulated at the 5΄end of the transcript. Interestingly, in the case of MERS-CoV Nsp1, we observed that host gene expression inhibition relies explicitly on translation inhibition and not mRNA degradation. Concerning previously suggested Nsp1 inhibitors, we found that Montelukast and Ametantrone do not affect the capacity of Nsp1 to inhibit host mRNA translation. Last, we show that Nsp1 from SARS-CoV-2, MERS, and Bat-Hp coronaviruses specifically protects viral mRNAs at the 5’UTR, suggesting a co-evolutionary adaptation of these viral traits.

## Material and Methods

### Cell lines and cell culture

HEK293 Flip-In cells were grown in DMEM (+/+) medium (Gibco, Cat. No. 41966-029), supplemented with 10% Fetal Bovine Serum (SIGMA, Cat. No. F7524-500ML) and 100 units/ml Penicillin-Streptomycin (Bio-Concept, Cat. No. 4-01F00-H) at 37°C and 5% CO_2_. The cells were detached with Trypsin-EDTA PBS (BioConcept, Cat. No. 5-51F00-H, 1:250). Cell counting was performed by Trypan Blue Stain (Gibco, Cat. No. 15250-061) using the automated LUNA II TM cell counter.

### Polysome profiling and Western blot analysis

75-90% confluent HEK293 cell were transfected in 150 mm dishes with either pcDNA 3.1(+) FLAG-SARS-CoV-2 Nsp1 WT or pcDNA 3.1 (+) FLAG-SARS-CoV-2-Nsp1-KH ^3^. The cells were treated with 4.25 μg of the corresponding plasmid using 7.5 μl μg^−1^ DNA Dogtor transfection reagent (Oz Biosciences, Cat. No. DT510000) in Opti-MEM medium (Gibco, Cat. No. 51985-026) overnight at 37°C and 5% CO_2_. For a positive ribosome collision control ^28^ 75-90% confluent HEK293 cells were cultured in 150 mm dishes and incubated with medium containing 1.8 µM Emetine (Millipore, Cat. No. 324693) for 15 minutes at 37°C and 5% CO_2_. After the medium supplemented with Emetine and the complete growth medium of the control condition were removed, the cells were washed twice with PBS (Gibco, Cat. No. 10010-031). The cells were then scraped with ice-cold PBS and transferred to a 1.5 ml microcentrifuge tube and further treated as described for the transfected cells.

15 and 50% sucrose gradient solutions were supplemented with 10 mM DTT (Merk, Cat. No. 43816-10ml) and 0.1 mM Cycloheximide in H_2_O (CAS. No. 66-81-9) and gradients were generated using a Biocamp Gradient Station ip in SW41 tubes (SETON, Cat. No. 7030).

Twenty-four hours post-transfection cells were treated with 100 μg/ml Cycloheximide in DMSO (CAS. No. 66-81-9) for 4 minutes at 37°C. The medium was removed, and the cells were harvested with PBS (Gibco, Cat. No. 10010-031) containing 100 μg/ml Cycloheximide (CAS. No. 66-81-9) by scraping and transferred into a 1.5 ml microcentrifuge tube. The cells were pelleted for 5 minutes at 4°C and 500 *g* and resuspended in 1x lysis buffer (10 mM Tris-HCl pH 7.5 (Fisher Bioreagent, Cat. No. BP1756-100), 10 mM NaCl (SIGMA, Cat. No. S5150-1L), 10 mM MgCl_2_ (Merck, Cat. No. A0136133 121), 1 mM Triton, 1mM Sodium deoxycholate (SIGMA-ALDRICH, Cat. No. D6750-256), 1 mM DTT (Merk, Cat. No. 43816-10ml), 0.1 mM Cycloheximide in DMSO (CAS. No. 66-81-9) and 0.2 U/μl Murine RNase inhibitor (Vayzme, Cat. No. R301-01-AA) (DTT, Cycloheximide and Murine RNase inhibitor were freshly added to the lysis buffer) and incubated for 2 minutes at 4°C. After incubation on ice, the cell lysate was centrifuged for 5 minutes at 16,000 *g*, the supernatant was collected and the amount of cells was adjusted to the smallest total RNA measurement using the Nanodrop measurements to ensure equal loading. 280 μl of lysed cells were loaded onto the gradients andhe tubes were spun in the SW 41 Ti rotor for 2 hours at 274’355 *g,* 4°C in a BECKMAN COULTER OptimaTM L-90K ultracentrifuge. The gradients were fractionated using the Biocamp PGF ip Pistion Gradient Fractionator at a speed of 0.3mm/s in 20 microcentrifuge tubes per replicate. During fractionation, the A_260_ profile was recorded and absorption measurements were visualized using Graph Pad Prism (version 10.0.2). The sucrose fractions were stored at - 80°C for further protein analysis.

The collected sucrose fractions were thawed on ice, mixed with 1 volume of acetone and 0.1 volume of 100% TCA (SIGMA-ALDRICH, Cat. No. T9159-2500) and incubated at -80°C for 4 hours, and then centrifuged at 16,000 *g* for 5 minutes at 4°C. The precipitated proteins were washed twice with 1ml of ice-cold acetone (centrifuged at 16,000 *g* for 5 minutes at 4°C) and pellets were dried for 20 minutes using the Eppendorf Concentrator plus and 45 μl 1.5x LDS loading buffer (Invitrogen, Cat. No. NP0008), containing 50 mM DTT (Merk, Cat. No. 43816-10ml), and 45 μl 8M urea (APOLLO Scientific Limited, Cat. No. BIU41110) were added and incubated for 5 minutes at 95°C. After heating, the samples were briefly vortexed. A 1:10 and a 1:5 dilution were prepared for the early fractions (free fractions) of each replicate.

Of each selected fraction, 8 μl was loaded into a NuPAGE 4-12% Bis-Tris (26-well) (Invitrogen Cat. No. WG1403A) or 20 μl into a mPAGE 4-12% Bis-Tris 15-well gel (Milipore, Cat. No. MP42G15). Gels ran for 35 to 40 minutes in MES-buffer (50 mM MES (SIGMA, Cat. No. M3671-250G), 50 mM Tris base (Fisher Bioreagents, Cat. No. BP152.5), 0.1% SDS (Fisher Bioreagents, Cat. No. BP2436-1), 1 mM EDTA (SIGMA-ALDRICH, Cat. No. E5134-500G), pH 7.3). The proteins were then transferred onto a nitrocellulose membrane using Trans-Blot Turbo Transfer Pack (BIO-RAD, Cat. No. 1704158). Ponceau S was used to check the completeness of the protein transfer. The membranes were then cut according to the mass of the proteins to be analyzed (RPL4: 50 kDa, FLAG-Nsp1: 25 kDa, EDF1: 16-20 kDa) and blocked in 5% milk in TBS-tween (0.1%) for 30 minutes and were incubated overnight at 4 °C under gentle agitation with Rabbit anti-RPL4 (Proteintech, Cat. No. 11302-1-AP), Rabbit anti-EDF1 (Abcam, Cat. No. ab174651), Mouse anti-FLAG (SIGMA-ALDRICH, Cat. No. 11302-1-AP), Rabbit anti-EDF1 (Abcam, Cat. No. ab174651), Mouse anti-FLAG (SIGMA-ALDRICH, Cat. No. SLBJ4607V). Membranes were washed 3x with TBS-tween (0.1%) for 5 minutes each time and incubated with Donkey anti-mouse 700CW (LI-COR, Cat. No. 926-68022) and Donkey anti-rabbit 800CW (LI-COR, Cat. No. 926-32212) as secondary antibodies for one hour at 4°C with agitation. The membranes were washed with TBS-tween (0.1%) once for 10 minutes and twice for 5 minutes before measuring the signal with the Odyssey Infrared Image System. The resulting images were labelled using Inkscape (version 1.2.2).

### Preparation of HeLa translation-competent lysate

Lysates were prepared from HeLa S3 cell cultures like previously ^29^ grown to a density ranging from 1 to 2 × 10^6^ cells/ml, pelleted at 200 *g*, 4°C for 5 min, washed twice with cold 1x PBS (pH 7.4) and resuspended in a translation-competent buffer, (33.78 mM HEPES (pH 7.3), 63.06 mM KOAc, 0.68 mM MgCl2, 54.05 mM KCl, 13.51 mM creatine phosphate, 229.73 ng/ml creatine kinase and 1x protease inhibitor cocktail (Bimake, Cat. No. B14002)) to a final concentration of 2×10^8^ cells/ml in 2 ml screw cap microtubes (Sarstedt, Cat. No. 72.693). The cells were lysed by dual centrifugation at 500 RPM at -5°C for 4 min using the ZentriMix 380 R system (Hettich AG) with a 3206 rotor with 3209 adapters. After dual centrifugation, the lysate was pelleted at 13,000 *g*, 4°C for 10 min. The supernatant (the lysate) was aliquoted, snap-frozen and stored at -80°C. The lysate could be thawed and refrozen multiple times for subsequent applications with only a minor loss of translation efficiency as reported in ^29^.

### Recombinant Proteins

All recombinantly expressed proteins of SARS-CoV-2, MERS-CoV and Bat-Hp-CoV Nsp1, as well as the control protein Ecoli-Trx-SARS-CoV-2-Nsp1-CTD KH-AA were produced as described previously ^9^. The recombinant proteins used for *in vitro* translation were dialyzed at a concentration of 25 µM overnight at 4°C against 30 mM NaCl, 5 mM HEPES (pH 7.3) using a Slide-A-Lyzer Mini Dialysis devices (Thermo Scientific, 88400). The concentration was determined with Nanodrop. An equal amount of protein storage buffer was dialyzed in the same manner as a control to complement each translation reaction with the same amount of the dialyzed buffer.

### *In vitro* Translation Assays

HeLa S3 lysate at a final concentration of 1.0×10^8^ cells per ml was supplemented to 1 u/μl of Murine RNase Inhibitor (Vazyme, Cat. No. R301-03). Control reactions contained 250 µg/ml of Puromycin (Santa Cruz Biotechnology, Cat. No. sc-108071). As controls, parallelly purified Trx-SARS-CoV-2-CTD-KH Nsp1 (Ctr) was used instead of Nsp1 protein variants at the same final concentration. Unless otherwise stated, 12.5 μl *in vitro* translation reactions were performed for 50 min at 33°C. Control protein and Nsp1 proteins were supplemented in a concentration of 0.6 µM or titrated from 0 µM to 1 µM. Before adding the mRNA reporter, the translation mixes containing recombinant proteins were pre-incubated at 4° for 15 minutes, then at 33°C for 5 minutes. The *RLuc in* vitro transcribed and capped reporter mRNAs were incubated at 65°C for 5 minutes and then stored directly on ice (for non-viral mRNAs) or left to refold at room temperature for 5 minutes before being placed on ice (for viral 5’UTR-containing reporter mRNAs) until pipetted to the translation reactions at a final concentration of 5 fmols/μl. *In vitro* translation assays used for subsequent mRNA degradation analysis were performed in a total volume of 75 µl instead of 12.5 µl and for 90 min. Negative control reactions contained 250 µg/ml of puromycin or 7.5 µg/ml of rocaglamide. To monitor the protein synthesis output by the luciferase assay, 5 µl of each translation reaction was mixed with 5 µl 1x Renilla-Glo substrate in Renilla-Glo assay buffer and supplemented with 40 µl of H_2_O (Promega, Cat. No. E2720) on a white-bottom 96-well plate (Greiner, Cat. No. 655073). The luminescence was measured with the TECAN infinte M1000 Pro plate reader after 15 minutes of incubating the microplate at room temperature. Three luminescence measurements were taken from each biological replicate, which were later averaged using Excel (version 2403) and plotted using GraphPad (version 10.0.2).

The cytoplasmically enriched translation-competent lysate supplemented with purified SARS-CoV-2 Nsp1 WT was used to evaluate the potential Nsp1 inhibitors, Montelukast Sodium (CAS Number 151767-02-1, Merck, Cat. No. Y0001434) and Amentantrone (ASTA TECH, Cat. No. I11677), in a dose-dependent manner. A 10 mM stock, stored in DMSO, was diluted with RNase-free water (SIGMA, Cat. No. W4502-1L) in a dilution series to the following solutions: 5 μM, 10 μM, 25 μM, 50 μM, 75 μM, 100 μM, 500 μM, 1000 μM. After addition of the inhibitor, purified SARS-CoV-2 Nsp1 WT was pipetted into the reactions at a final concentration of 0.5 μM.

### Cloning, Expression and Purification of RLuc Reporter Plasmids

MS2-containing mRNA reporters with the viral leader sequence of SARS-CoV-2 WT (p200-T7mini+G-SARS-CoV-2-hRluc-6xMS2) were used to derive the mutant versions; scrambled SL1 (scrSL1), deletion of SL1 (ΔSL1), apical mutation of SL1 (ap. mut. SL1). PCR reactions contained 0.3 µM of each primer and 1 ng/µl of plasmid. The p200-T7mini+G-SARS-CoV-2-ap.mut.SL1-hRluc-6xMS2 plasmid was obtained after QuikChange® Site-Directed Mutagenesis with the mutagenic primer ATT AAA GGT TTA TAC CTA GGG AGG TAA CAA ACC AAC CAA CTT T. The other versions of SARS-CoV-2 reporter mRNA were cloned by In-Fusion® HD Cloning. The vectors were amplified by CloneAmp HiFi PCR Premix (TaKaRa, Cat. No. ST0506) from the p200-T7mini+G-SARS-CoV-2-hRluc-6xMS2 and with following primers; TAC GAC TCA CTA TAG AAC CAA CTT TCG ATC TCT TGT AGA TCT G, GAT CGA AAG TTG GTT CTA TAG TGA GTC GTA TTA CAA TTC ACT GGC C (ΔSL1), GGA TCG ATA CAT ATA ACG GCT TAA CTA ATC CAT, ATG GAT TAG TTA AGC CGT TAT ATG TAT CGA TCC (scrSL1). The primers accounted for the deletion of SL1 or contained the scrambled sequence of SL1 included respectively.The PCR products were treated with Cloning Enhancer (TaKaRa, Cat. No. ST0003) at 37°C for 15 min and subsequently at 80°C for another 15 min. The plasmids were re-ligated with 5x In-Fusion Enzyme Premix (TaKaRa, Cat. No. ST0345) at 50°C for 15 min. The In-Fusion reactions were transformed in home-made Stellar competent cells and plated on Agar plates with appropriate antibiotics. Correct colonies were confirmed by sequencing, and from liquid bacterial stocks, the plasmids were isolated using the ZymoPURE II Plasmid Midiprep Kit (Zymo Research, Cat. No. D4201-B) according to the manufacturer’s instruction. Viral reporter mRNAs were described previously^9^.

### *In vitro* Transcription of Reporter RLuc mRNA

DNA plasmids for *in vitro* transcription reactions are all based on a vector with a shortened T7 promoter to reduce the distance between the SL1 loop of the viral reporters and the cap as described in ^9^. All plasmids were linearized in reactions containing 40 ng/ml DNA and 0.5 U/ml restriction enzyme (HindIII-HF, NEB, Cat. No. R3104L) in 1x CutSmart buffer (NEB, Cat. No. B7204S). After confirming linearization by agarose gel electrophoresis the reactions were purified using the MACHEREY-NAGEL NucleoSpin Gel and PCR Clean-up kit (Cat. No. 740609.50) quantified by measuring OD260 with Nanodrop. The linearized plasmid served as a template in reactions containing 1x OPTIZYME transcription buffer (Thermo Fisher Scientific, Cat. No. BP81161), 1 mM of each ribonucleotide (Thermo Fisher Scientific, Cat. No. R0481), 1 U/ml RNase inhibitor (Vazyme, Cat. No. R301-03), 0.001 U/ml pyrophosphatase (Thermo Fisher Scientific, Cat. No. EF0221) and 1 U/ml T7 polymerase (Thermo Fisher Scientific, Cat. No. EP0111). After an incubation step of 1 h at 37°C an equal amount of polymerase was added to boost transcription for another 30 min. Subsequently, the template DNA was degraded with 0.14 U/ml Turbo DNase (Invitrogen, Cat. No. AM2238) for 30 min at 37°C. The transcribed reporter mRNA was isolated using the Monarch RNA Cleanup Kit (New England Biolabs, Cat. No. T2040L), eluted with nuclease-free water and quantified as mentioned above.

A 7-methylguanylate cap was added to the reporter mRNAs using the Vaccinia Capping System from New England Biolabs (Cat. No. M2080S) according to the manufacturer’s instructions with the addition of 1U/ml RNase inhibitor (Vazyme, Cat. No. R301-03). The resulting capped mRNA was purified, eluted, and quantified as described above. All reporter mRNAs contain an 80-nt nucleotide-long template-encoded poly(A) tail.

### Reverse Transcription and qPCR

The RNA from *in vitro* translation reactions was isolated using the Maxwell RSC simplyRNA Cells kit (Promega, Cat. No. AS1390) with the Maxwell RSC Instrument following the manufacturer’s guidelines. The RNA was eluted in kit-provided ribonuclease-free water, and concentrations were quantified using nanodrop. All RNA isolates were diluted to 100 ng/µl of total RNA.

Reverse transcription reactions contained 500 ng of total RNA, 9 ng/ µl of random hexamers (Microsynth), 1x AffinityScript RT buffer (Agilent, Cat. No. 600100-52), 10 mM DTT, 0.4 mM dNTP mix (each ) (Vazyme, Cat. No. P031-01), 0.4 U/μl Murine RNase inhibitor (Vazyme, Cat. No. R301-03) and 2% AffinityScript Multiple Temperature Reverse Transcriptase (Agilent, Cat. No. 600100-51). After incubation of 5 min at 65°C of RNA with random hexamers, the reaction was cooled at room temperature for 10 min before adding RT+ or RT-master mix. The RT-master mix lacked the Reverse Transcriptase as negative control. The cDNA synthesis was performed at 50°C for 1 hour before inactivating the enzyme at 75°C for 15 min. Finally, the cDNA was diluted to 1 ng/µl for consequent qPCR analysis.

For qPCR, each sample contained 0.2 ng/µl of cDNA and 0.5 µM of forward and reverse primer. As a normalizer, the mRNA levels of GAPDH were monitored with the primer set, GAG TCA ACG GAT TTG GTC G and GAG GTC AAT GAA GGG GTC AT. mRNA levels of the non-viral RLuc reporter were observed in the 5 ‘UTR with the oligonucleotides TCT GCA GAA TTC GCC CTT CAT G and GCA CGT TCA TTT GCT TGC A. For the 3’ UTR of the mRNA two primers were used: GAG AAG GGC GAG GTT AGA CG and TGG AAA AGA ACC CAG GGT CG. The viral 5’ UTR reporter of SARS-CoV-2 was monitored with the primer pair: GGT TTA TAC CTT CCC AGG TAA CAA ACC and GCC GAG TGA CAG CCA CAC AG. The quantitative PCR was executed with 1x Brilliant III Ultra-Fast SYBR Green QPCR Master mix (Agilent, Cat. No. 600882) in the Rotor-Gene 6200 real-time system. The fluorescence readout and cycle values (CT-values), with a manually set threshold at 0.1, was quantified using the Rotor-Gene 6200 software. Subsequently, the relative mRNA levels were calculated using the comparative CT method.

### Northern Blot Analysis

Probes specific for the target RNAs were synthesized by PCR with 1x Clone Amp HiFI Premix (TaKaRa, Cat. No. St0506), 0.04 ng/µl of plasmid and 0.3 µM of each forward and reverse primer. The oligonucleotides TCT GCA GAA TTC GCC CTT CAT G and GCA CGT TCA TTT GCT TGC A were used to amplify the 5’ UTR region of the non-viral reporter mRNA (p200-T7mini+G-hRluc-6xMS2), and GAG AAG GGC GAG GTT AGA CG and TGG AAA AGA ACC CAG GGT CG for the 3’ end respectively. GGT TTA TAC CTT CCC AGG TAA CAA ACC and GCC GAG TGA CAG CCA CAC AG were employed to produce the probe annealing in the 5’ UTR of SARS-CoV-2. The PCR products were isolated and purified by the Gel clean up guidelines of the NucleoSpin Gel and PCR clean-up Kit (Macherey-Nagel, Ref. No. 740609.50).

For Northern blot analysis, 500 ng of total isolated RNA from the in vitro translation reactions was denatured in NorthernMax Formaldehyde Load Dye (Thermo Scientific, Cat. No. AM8552) supplemented with EtBr (10 µg/ml) at 65°C for 15 min. Electrophoresis in 1x RNase-Free MOPS buffer (200mM MOPS (pH 7.0), 80mM Sodium acetate, 10mM EDTA ) at 75 V (max 200 mA) for 3.5 hours at room temperature resolved the RNA on a 1.2% agarose gel containing 1% (w/v) formaldehyde and 1x MOPS buffer. The gel was transferred to a positively charged Amersham Hybond N+ membrane (Cytiva, Cat. No. RPN303B) overnight using capillary action with 20x SSC buffer (Sigma-Aldrich, Cat. No. 33220). Following transfer, the RNA was cross-linked to the membrane by UV irradiation in a Stratalinker UV irradiation chamber at 0.12 J for 1 min. A brief staining with 0.03% (w/v) methylene blue dye, 300 mM sodium acetate visualized 28 S and 18 S rRNA which is used as loading control. After staining, the membrane is destained with Milli-Q water and dried. The dried membranes were prehybridized with ULTRAhyb Ultrasensitive Hybridization Buffer (Thermo Scientific, Cat. No. AM8670) at 42°C for 30 min to minimize nonspecific binding. The PCR probes were labelled with α^32^P-dCTP using the DecaLabel DNA Labelling Kit (Thermo Scientific, Cat. No. K0622) and cleaned up using Amersham MicroSpin G-25 Columns (Cytiva, Cat. No. 27532501). The labelled supplemented to hybridization buffer at a final concentration of 10 ng/ml was incubated overnight at 42°C with the membranes. After completion of hybridization, the membranes were washed sequentially with low-stringency wash buffer (2x SSC, 0.1% SDS) two times for 5 min and high-stringency wash buffer (0.1x SSC, 0.1% SDS) two times for 15 min at 42°C to remove non-specific binding of the probes. Finally, the membranes were exposed to a phosphor screen overnight, and the radioactivity signal was visualized using the Typhoon FLA 9500 detection system at 1000V 100 μm. The hybridized bands were quantified by ImageJ and normalized to the control protein-containing reactions.

## Results

### Nsp1 promotes mRNA degradation independently of ribosome collision

Ribosome collisions during translation are a sign of malfunction and can lead to the activation of quality control pathways to maintain cellular homeostasis ^30^. Collided ribosomes are sensed and stimulate ribosome-associated quality control (RQC) pathways, potentially leading to the degradation of mRNA and nascent peptides ^24,31^. Given that Nsp1 binds the ribosome and stimulates mRNA degradation, we set out to assess whether Nsp1 stimulates global ribosome collisions.

We expressed N-terminally 3×FLAG-tagged Nsp1 of SARS-CoV-2 WT or the mutant KH-AA (positions 164-165) that cannot bind the ribosome in HEK293T cells. To monitor ribosome collision occurrence, we performed polysome profiling to assess whether EDF1, which acts as a ribosome collision sensor and whose conserved binding site is located in the 40S subunit, associates with polysomes upon Nsp1 expression ^28^. As expected, the KH-AA mutant localized solely to the free fractions, whereas the WT Nsp1 localized to the 40S fraction, as previously reported ^2^. Furthermore, the expression of WT Nsp1 leads to an increase in the 80S monosome peak and a decrease in the polysome signals compared to the expression of the KH-AA mutant Nsp1 (Fig.1A), as observed previously ^2^. In both cases, EDF1 was found primarily in the free fraction and bound to 40S ribosomes, indicating no occurrence of 80S ribosome collisions (Fig. 1). As a positive control experiment for ribosome collisions, we incubated cells with a low dose of emetine (1.5 μM for 15 min) ^28^. The treatment led to the accumulation of EDF1 in the polysomal fractions (Sup. Fig. 1).

**Figure 1.**
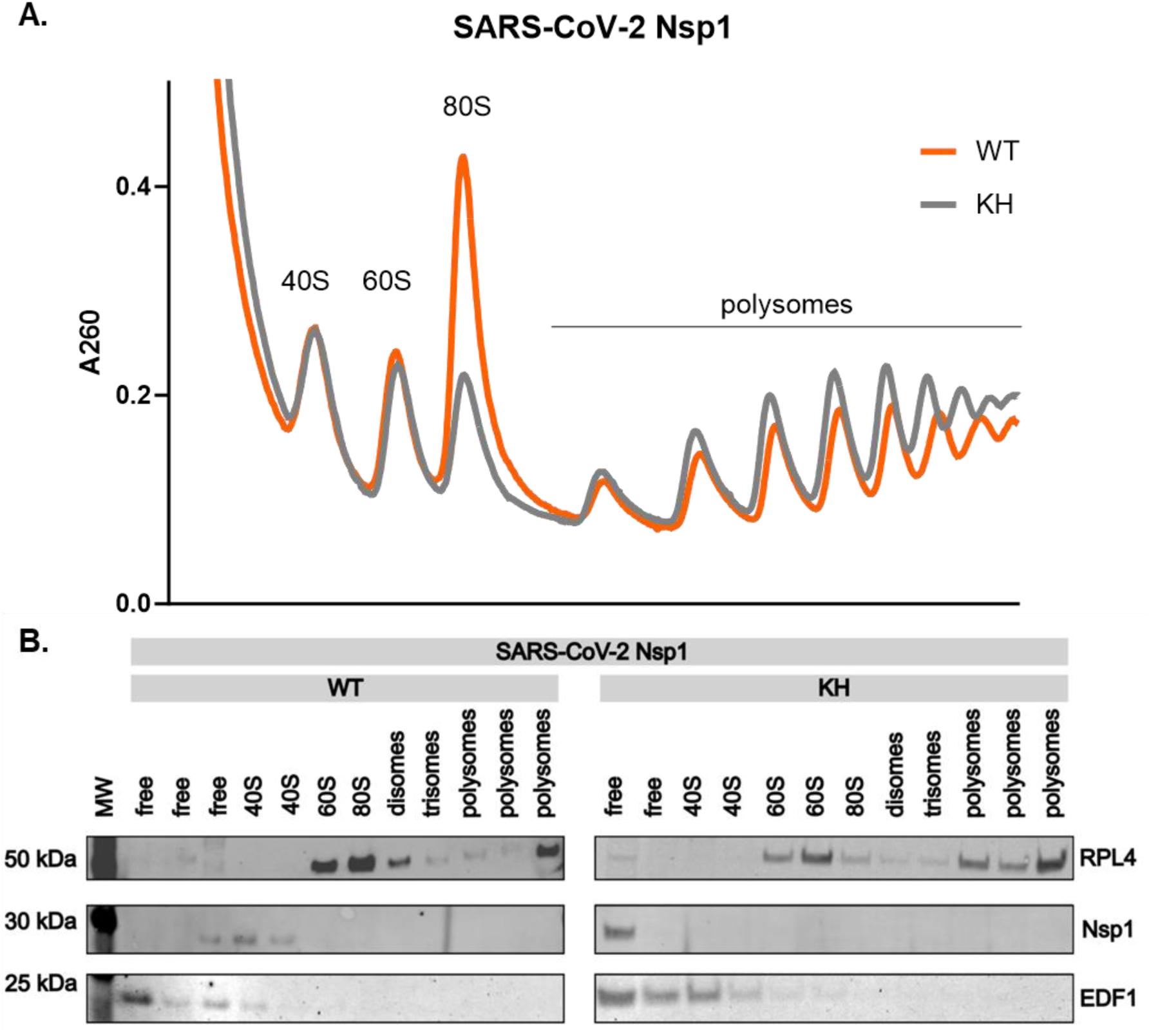
Nsp1-dependent mRNA degradation is not stimulated by ribosome collisions. **(A)** Equal amounts of cell lysates from HEK293T cells 24h after transfection with plasmids coding for FLAG-SARS-CoV-2 Nsp1 WT (WT) or FLAG-SARS-CoV-2 Nsp1 KH (KH) were fractionated on a 15-50% sucrose gradient by ultracentrifugation. **(B)** From the collected polysome fractions, total proteins were extracted with acetone and TCA. Detected proteins are labelled on the right side of the blots.

Our data suggests that Nsp1-dependent host mRNA degradation occurs independently of generic ribosome collision-induced quality control pathways.

### Nsp1 binding to host ribosomes is required for host mRNA degradation

The fact that Nsp1 does not lead to ribosome collisions urged us to assess whether Nsp1-mediated mRNA degradation requires active translation or if binding of Nsp1 to the 40S subunit suffices to degrade host cell mRNAs.

We investigated whether Nsp1 translation inhibition is accompanied by RNA degradation using a human-based *in vitro* translation system produced from dual-centrifugation-treated HeLa S3 cells ^29^. This way, we monitor inhibition within a shorter time frame than experiments performed in living cells. We performed *in vitro* translation of capped and polyadenylated *Renilla* Luciferase (Rluc) reporter mRNA with a non-viral 5΄UTR in the presence of SARS-CoV-2 Nsp1 protein variants (Fig. 2A). We monitored the translation signal by luciferase assay and the RLuc reporter mRNA levels by Northern blot analysis and RT-qPCR.

**Figure 2.**
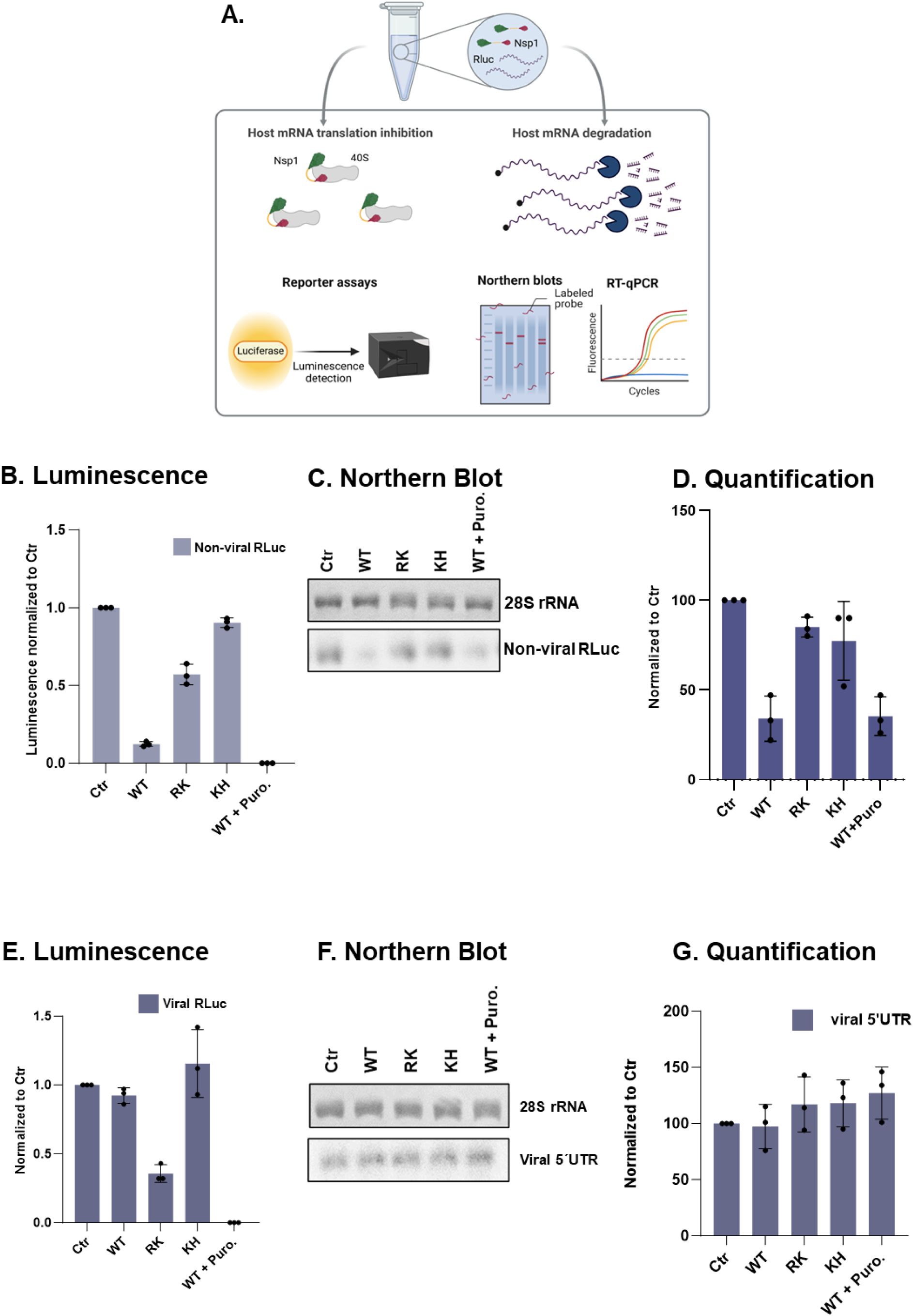
Decoupling the translation inhibition and degradation activities of SARS-CoV-2 Nsp1 using a human-based cell-free translation system. **(A)** Schematic overview of the *in vitro* translation assay followed by RLuc measurements to monitor translation efficiency and Northern blot and qPCR to assess mRNA stability in the presence of Nsp1 variants. **(B)** Relative RLuc activity measurements of 50 min *in vitro* translation reactions with 0.6 µM of purified SARS-CoV-2 Nsp1 WT, RK-AA or KH-AA mutant, normalized to the reaction containing the purified control protein (Ctr) after three measurements. **(C)** 500 ng of isolated RNA from the *in vitro* translation reactions was subjected to Northern blot analysis using a ^32^P-labelled probe for the reporter RLuc RNA. The methylene blue-stained pattern of the 28S rRNA is shown as an internal standard for RNA loadings for each lane. **(D)** Northern blot quantification of C **(E)** Relative RLuc activity measurements as in (B) using an RLuc reporter mRNA with the full-length SARS-CoV-2 5΄UTR. **(F)** Northern blot analysis as in (C) using the reporter of panel (E). **(G)** Northern blot quantification of C. Data in panels (B) and (E) are presented as mean values of three biological replicates (sets of translation reactions) averaged after three measurements. Samples of the same translation reactions were then used for northern blot and qPCR analysis.

Apart from WT Nsp1, we tested the ribosome-binding mutant of Nsp1 (KH-AA), which abolishes the interaction of the C-terminal end of the protein with the ribosomal entry channel ^2,3^. A second mutant, RK-AA (positions 124-125), was also analyzed. This mutant cannot induce host-cell mRNA degradation, exhibits reduced efficiency in inhibiting mRNA translation, and fails to protect viral 5′ UTRs from Nsp1-mediated mRNA translation inhibition and degradation ^9,11^. We translated *in vitro* transcribed and capped reporter mRNAs with equal amounts of purified Nsp1 proteins. We normalize the impact of recombinant Nsp1 proteins using a control (Ctr) protein, which consists of bacterial thioredoxin fused to the KH-AA mutant form of Nsp1 at its C-terminal end. This fusion protein is of similar size to wild-type Nsp1. The use of this Ctr protein allows us to rule out any effects stemming from potential contaminants introduced during protein purification, as it mimics the size and composition of Nsp1 without retaining its functional activity.

As reported previously, the RLuc readout showed an apparent reduction of translation in the presence of WT Nsp1 protein (Fig. 2B). The presence of the KH-AA Nsp1 mutant abolished the capacity of Nsp1 to inhibit translation, and the RK-AA mutant led to a half-way decrease of the RLuc output compared to the WT condition.

To determine if Nsp1 stimulates mRNA degradation in our lysate, we purified and analyzed mRNA integrity by Northern blot after *in vitro* translation. Northern blot analysis revealed a considerable reduction in the full-length mRNA level in the presence of WT Nsp1 (Fig. 2C-D). Conversely, the mRNA remained practically unaffected in the presence of Nsp1 RK-AA. Notably, we observed that in translation reactions containing WT Nsp1 and puromycin, a translation elongation inhibitor, reporter mRNAs were degraded to a similar extent as in the absence of puromycin. This result suggests that Nsp1 can induce mRNA degradation even under translation elongation inhibition.

We repeated the *in vitro* translation experiments followed by Northern blot analysis with an RLuc reporter mRNA harbouring the full-length SARS-CoV-2 5΄UTR that is otherwise identical in its features with the previous non-viral reporter mRNA. Similarly to previous reports from Nsp1 transfection experiments and viral infection experiments, cell-free translation recapitulated the protection effect of the viral 5΄UTR against Nsp1-mediated inhibition (Fig.2E-G). With RK-AA Nsp1, we observed a reduction in the ability of the protein to protect the reporter mRNA from translation inhibition (Fig.2E-G).

These results suggest that our *in vitro* translation system can recapitulate well-reported SARS-CoV-2 Nsp1 features. The RK Nsp1 mutant still inhibits mRNA translation without causing degradation, showing that our system can decouple the two processes. Moreover, the incubation of the lysate with Nsp1 and puromycin shows that elongation is not a prerequisite for Nsp1-mediated mRNA degradation.

### Nsp1 mediates mRNA Decay close to the 5’ End of Non-viral mRNAs

There is evidence that Nsp1-mediated mRNA degradation occurs within the 5΄UTR of the host cell mRNAs ^16^. To assess whether this is the case in our system, we performed a time-course experiment for 0, 25, 50 or 75 minutes of translation, incubating lysates with either a CTR protein or WT Nsp1. We analyzed the output of the *in vitro* translation reactions by monitoring the RLuc signal, which yielded the expected reduction of the RLuc protein output in the presence of WT Nsp1 (Fig. 3A). To assess the stability of the reporter RLuc mRNA, we used 5΄and 3΄-specific probes to determine whether degradation is more pronounced in the 5΄ end of the transcript (Fig. 3B). Northern blot experiments showed an overall degradation of the reporter mRNA over time even in the presence of the control protein. WT Nsp1 caused a pronounced time-dependent decay of the reporter mRNA towards the 5΄end, whereas the signal remained more stable when we used a probe that binds a sequence towards the 3΄end (Fig. 3C-D). This result suggests that cleavage occurs close to the 5΄ end of the transcripts because the signals of full-length molecules persist when using a 3΄-specific probe. In all cases, no additional degradation was observed in the presence of a control protein nor in the presence of the ribosome binding Nsp1 mutant (KH-AA), highlighting again that Nsp1 needs to bind the 40S to induce degradation. Interestingly, WT Nsp1 inhibits translation at the time point of 25΄ incubation when the mRNA transcript is still intact (compared to the sample containing a CTR protein of 25΄). This observation indicates that Nsp1-induced translation inhibition contributes more to the decreased protein output than the degradation of the reporter mRNA. This result demonstrates that we can decouple mRNA translation inhibition from the Nsp1-induced mRNA degradation in this condition.

**Figure 3.**
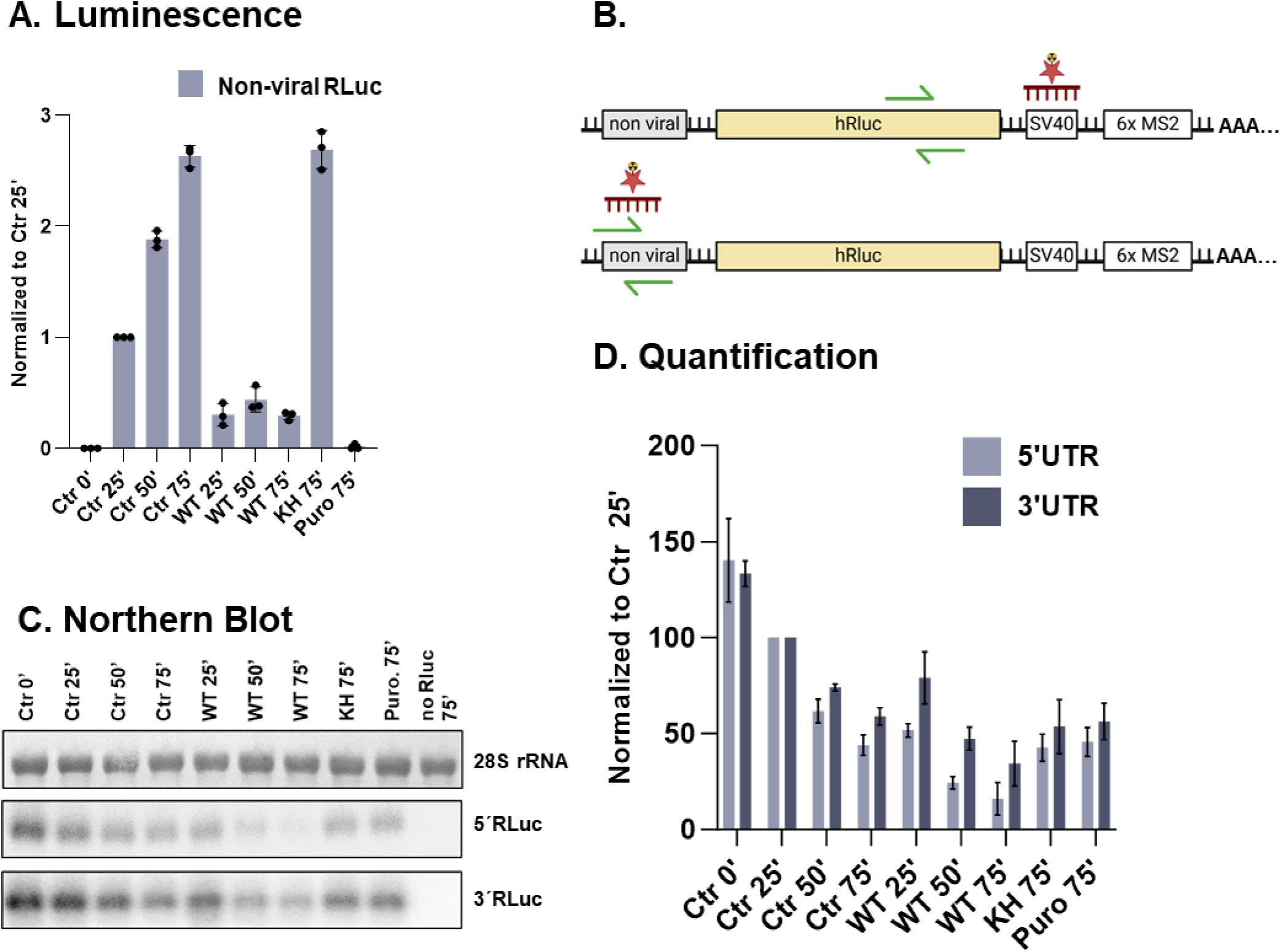
Nsp1 induces mRNA decay in the 5’ End of non-viral mRNAs. **(A)** Relative RLuc activity measurements of *in vitro* translation reactions with 0.6 μM of purified SARS-CoV-2 Nsp1 WT for increasing time normalized to the reaction containing the control protein (Ctr) (25 min of translation). **(B)** Schematic representation of the ^32^P-labelled probes and qPCR primers, annealing either in the 5΄UTR or 3΄UTR of the reporter mRNA. **(C)** Northern blot analysis of *in vitro* translation reactions described in A. A total of 500 ng isolated RNA from the *in vitro* translation reactions was subjected to northern blot analysis using a ^32^P-labelled probe binding the 5’-UTR or 3’UTR of the reporter RLuc RNA. The methylene blue-stained pattern of the 28S rRNA is shown as RNA loading control for each lane. **(D)** Northern blot quantification of C.

### Nsp1 promotes mRNA degradation even when translation is fully arrested

Intrigued by the finding that Nsp1 still stimulates mRNA degradation in the presence of puromycin (Fig. 2), we set out to explore whether steps of mRNA translation are a prerequisite for Nsp1 to stimulate mRNA degradation. To address this question and to exclude that mRNA degradation requires ribosomal scanning, we used the ribosome scanning inhibitor rocaglamide ^33^. We performed *in vitro* translation reactions in the presence of rocaglamide or puromycin with WT or KH Nsp1 and monitored the translation output by luciferase assay and the stability of the reporter mRNA using a 5΄ UTR-specific probe. We observed that WT Nsp1 also led to reporter mRNA degradation in the presence of rocaglamide, suggesting that Nsp1 can stimulate degradation without scanning (Fig.4A-C).

**Figure 4.**
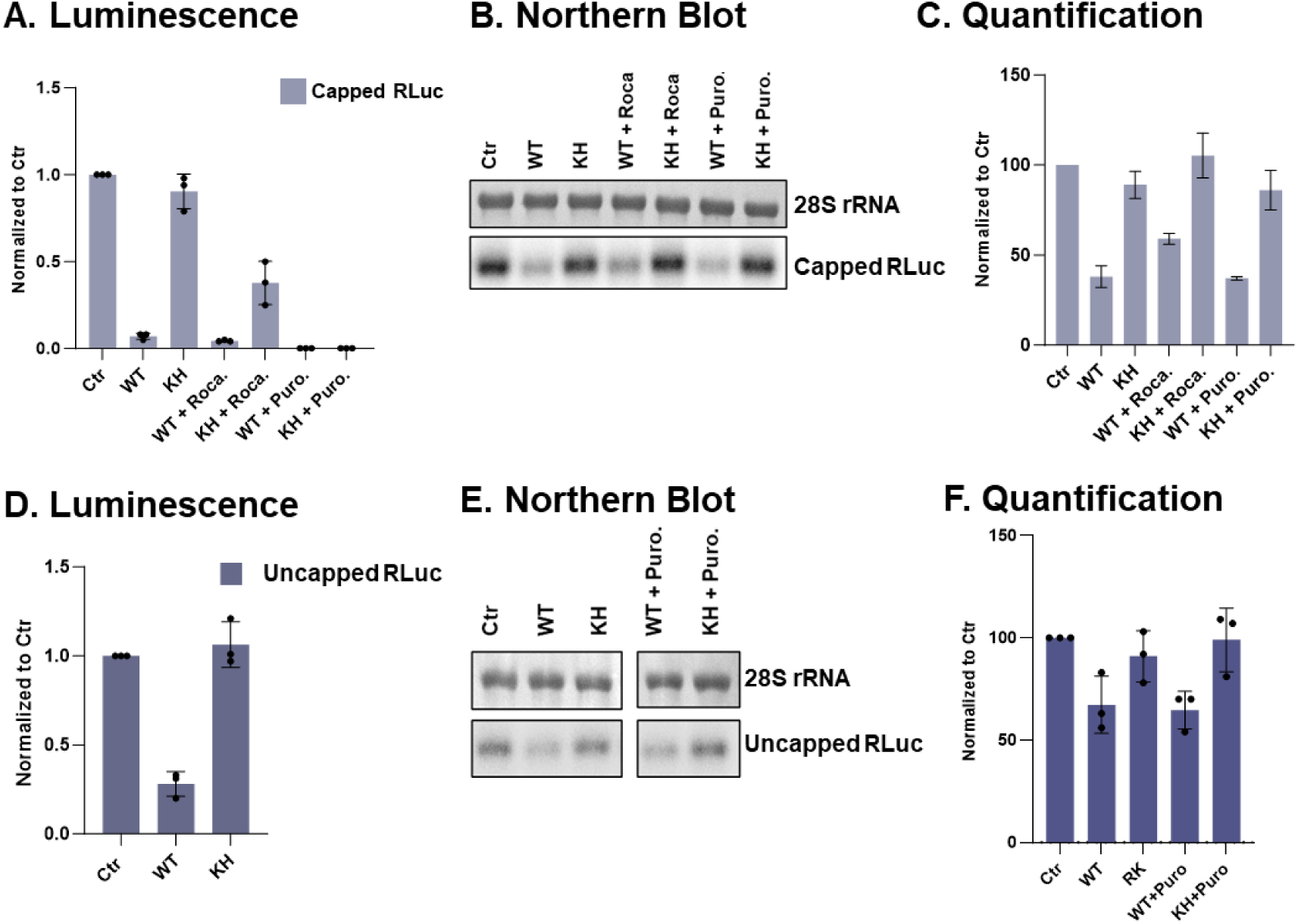
Nsp1-mediated RNA degradation does not require active translation. **(A)** Relative RLuc activity measurements of 50 min *in vitro* translation reactions with 0.6 µM of purified SARS-CoV-2 Nsp1 WT or KH mutant and capped non-viral mRNA, supplemented with 7.5 µg/ml of rocaglamide or 250 µg/ml of puromycin, normalized to the reaction containing the control protein (Ctr). **(B)** 500 ng isolated RNA from the *in vitro* translation reactions were subjected to Northern blot analysis using a ^32^P-labelled probe for the 5’-UTR of the reporter RLuc RNA. The methylene blue-stained pattern of the 28S rRNA is shown as an internal standard for RNA loadings for each lane. **(C)** Northern blot quantification of B. **(D)** Relative RLuc activity measurements of *in vitro* translation reactions with 0.6 µM of purified SARS-CoV-2 Nsp1 WT or KH mutant and uncapped non-viral mRNA, with 7.5 µg/ml of rocaglamide or 250 µg/ml of puromycin, normalized to the reaction containing the control protein (Ctr). **(E)** 500 ng isolated RNA from the *in vitro* translation reactions were subjected to northern blot analysis using a ^32^P-labelled probe for the 5’-UTR of the reporter RLuc RNA. The methylene blue-stained pattern of the 28S rRNA is shown as an internal standard for RNA loadings for each lane. Data in panels (A) and (C) are presented as mean values of three biological replicates (sets of translation reactions) averaged after three measurements. Samples of the same translation reactions were then used for northern blot analysis. **(F)** Northern blot quantification of E.

To verify that Nsp1 induces mRNA degradation independently of active translation, and because rocaglamide did not lead to complete translation inhibition, we assessed the impact of Nsp1 using uncapped mRNAs. Nsp1 still inhibited mRNA translation in a ribosome-binding-dependent manner (Fig. 4D). Northern blot analysis of the derived mRNAs revealed that degradation ensued in the absence of cap with and without puromycin (Fig. 4E-F). This evidence supports that neither canonical initiation nor translation elongation is required for Nsp1-induced mRNA degradation.

### MERS-CoV Nsp1 can inhibit mRNA translation independently of mRNA degradation

In the case of SARS-CoV-2 Nsp1, the inhibition of host gene expression is based, at least in part, on the degradation of host mRNA (reviewed in ^1^). This activity is in agreement with our human-based cell-free translation system results. We and others recently showed that MERS-CoV Nsp1 acts in a similar manner to inhibit host mRNA translation by binding through its C-terminal end to the small ribosomal subunit and blocking the mRNA entry channel ^8,9^. To test whether MERS Nsp1 can stimulate non-viral mRNA degradation, we incubated MERS-CoV WT Nsp1 or a mutant that cannot bind the 40S (KY-AA, positions 181-182) in human translation-competent lysates followed by *in vitro* translation of a RLuc reporter mRNA. As previously reported, we observed a robust translation inhibition in the presence of WT MERS-CoV Nsp1 that was abolished when the ribosome-binding mutant KY-AA was used (Fig. 5A).

**Figure 5.**
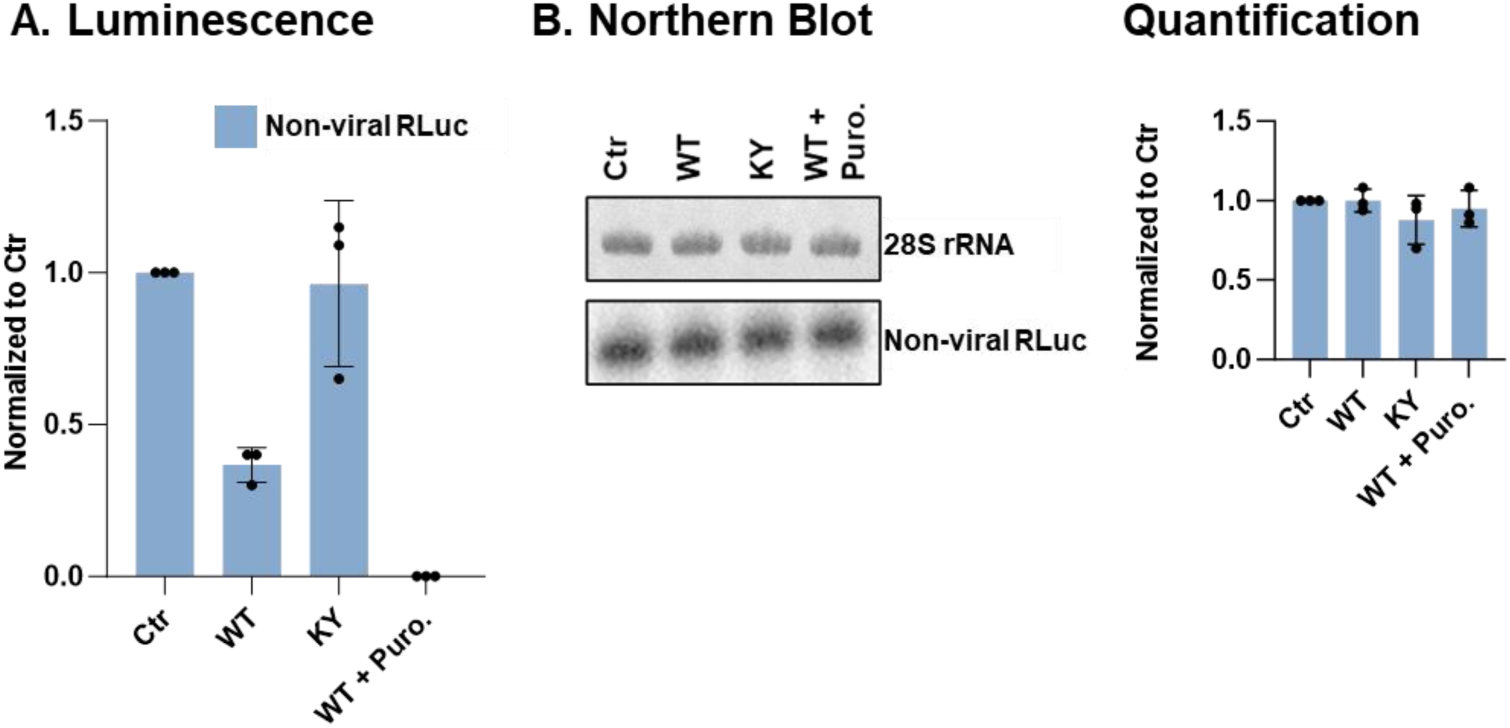
MERS-CoV Nsp1 can inhibit mRNA translation independently of mRNA degradation. **(A)** Relative RLuc measurements of *in vitro* translation reactions in the presence of MERS-CoV WT Nsp1, normalized to the control reaction containing a Ctr protein. Data are presented as mean values of three biological replicates (sets of translation reactions) averaged after three measurements. Samples of these translation reactions were then used for northern blot analysis in (B). **(B)** 500 ng isolated RNA from the *in vitro* translation reactions were subjected o northern blot analysis using a ^32^P-labelled probe for the 5’-UTR of the reporter RLuc RNA. The methylene blue-stained pattern of the 28S rRNA is shown as an internal standard for RNA loadings for each lane. Quantification of Northern blot signals from three biological replicates.

However, subsequent assessment of the RLuc mRNA reporter stability after translation by Northern blot and qPCR does not reveal hints of degradation in the sample containing MERS-CoV Nsp1 WT (Fig. 5B). These results signify that in our system, MERS-CoV Nsp1 can stimulate non-viral mRNA translation inhibition without affecting its stability, indicating differences in the mode of action of Nsp1 between SARS-CoV-2 and MERS-CoV.

### Assessment of potential Nsp1 inhibitors

Having established the means to uncouple Nsp1-mediated translation inhibition from mRNA degradation, we employed our cell-free translation system to assess whether previously reported small molecules inhibit the activity of Nsp1. The anthracenedione compound ametantrone has been reported to inhibit SARS-CoV-2 Nsp1-mediated RNA degradation ^21^. To test the effect of ametantrone on mRNA stability in our system, we supplemented HeLa S3 translation reactions with an RLuc reporter mRNA and purified WT SARS-CoV-2 Nsp1. The protein synthesis output at different concentrations of ametantrone was then tested using a luciferase assay. The stably decreased signal in the presence of varying ametantrone concentrations of the potential inhibitor showed that ametantrone did not inhibit the capacity of the SARS-CoV-2 Nsp1 to inhibit translation at any of the concentrations tested (Fig. 6A).

**Figure 6.**
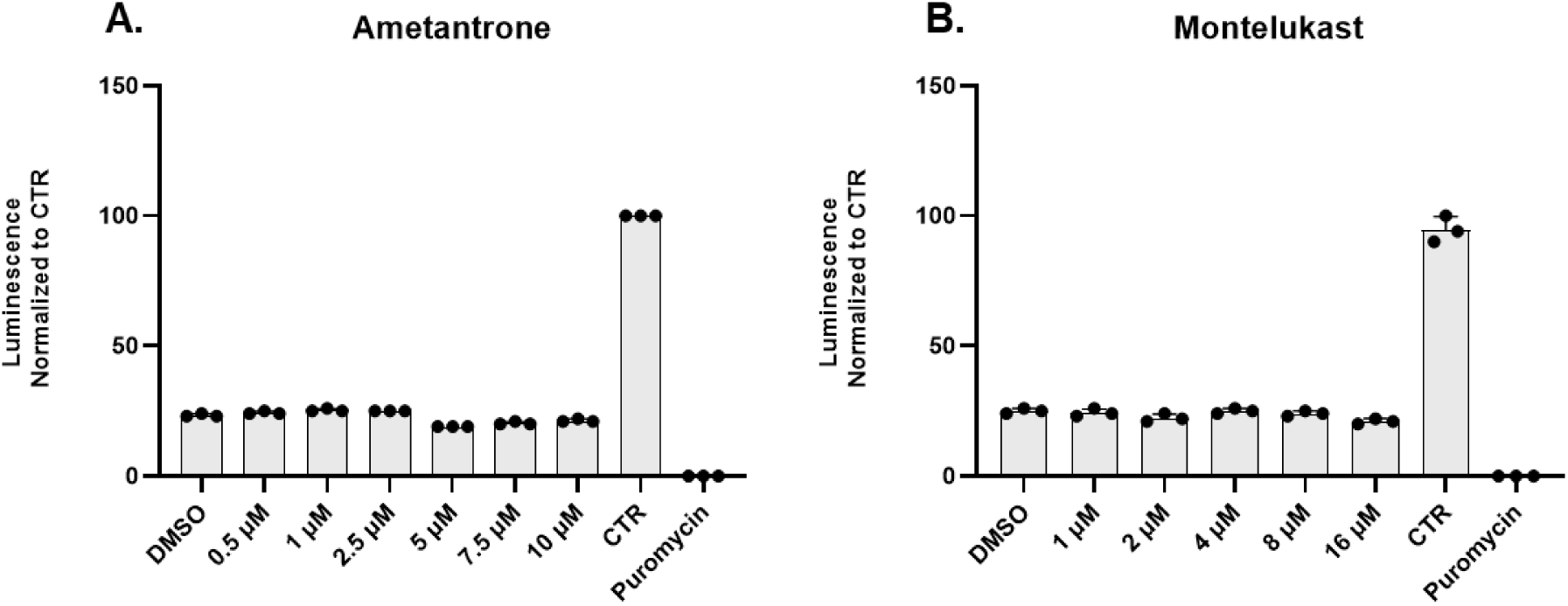
Assessment of potential SARS-CoV-2 Nsp1 inhibitors. Relative *RLuc* luminescence measurements of *in vitro* translation reactions with increasing concentrations of Ametantrone **(A)** or Montelukast **(B)** in the presence of SARS-CoV-2 Nsp1 WT, normalized to the reaction in the presence of the control protein (CTR) and no drug. Data are presented as mean values of three biological replicates (sets of translation reactions) averaged after three measurements. The positive control reaction for a translation inhibition contained 1 mg/mL puromycin.

An *in silico* study suggested that Montelukast sodium hydrate (an FDA-approved leukotriene receptor antagonist for asthma) can bind to the C-terminal of Nsp1 ^25^, rendering Montelukast a promising Nsp1 inhibitor. Similarly to Ametantrone, Incubation of Nsp1-containing lysates with different concentrations of Montelukast did not affect the capacity of Nsp1 to inhibit host cell translation (Fig. 6B), highlighting that the quest for potential Nsp1 inhibitors is still open. Notably, neither of the two inhibitors could inhibit translation when they were tested without Nsp1 (Sup. Fig. 8)

### Nsp1 portrays species-specific protection from translation inhibition

Ample evidence supports that SARS-CoV-2-encoded RNAs are not inhibited by Nsp1 ^9,10,34,35^ due to the presence of a 72-nt leader sequence at the very 5’ terminus, allowing them to bypass Nsp1-mediated translation inhibition. A sequence included in the stem-loop 1 (SL1), a *cis-*acting element encoded in all viral RNAs, is sufficient for evasion of Nsp1 inhibition ^5,10,11,34–36^. Interestingly, SL1 is the most variable region of the 5΄UTRs from different β-Coronaviruses, implying a possible co-evolution with the corresponding Nsp1 proteins ^37^. To address whether our *in vitro* translation system can recapitulate SL1 features that are important to evade Nsp1-mediated translation inhibition, we prepared RLuc reporter mRNAs harbouring the SARS-CoV-2 WT 5΄UTR, a variant with a scrambled SL1 and another lacking SL1 and a variant with a 4-nucleotide substitution of the apical sequence of SL1 that has been reported to be crucial for Nsp1-mediated protection from translation inhibition (Fig. 7A, upper panel) ^23^.

**Figure 7.**
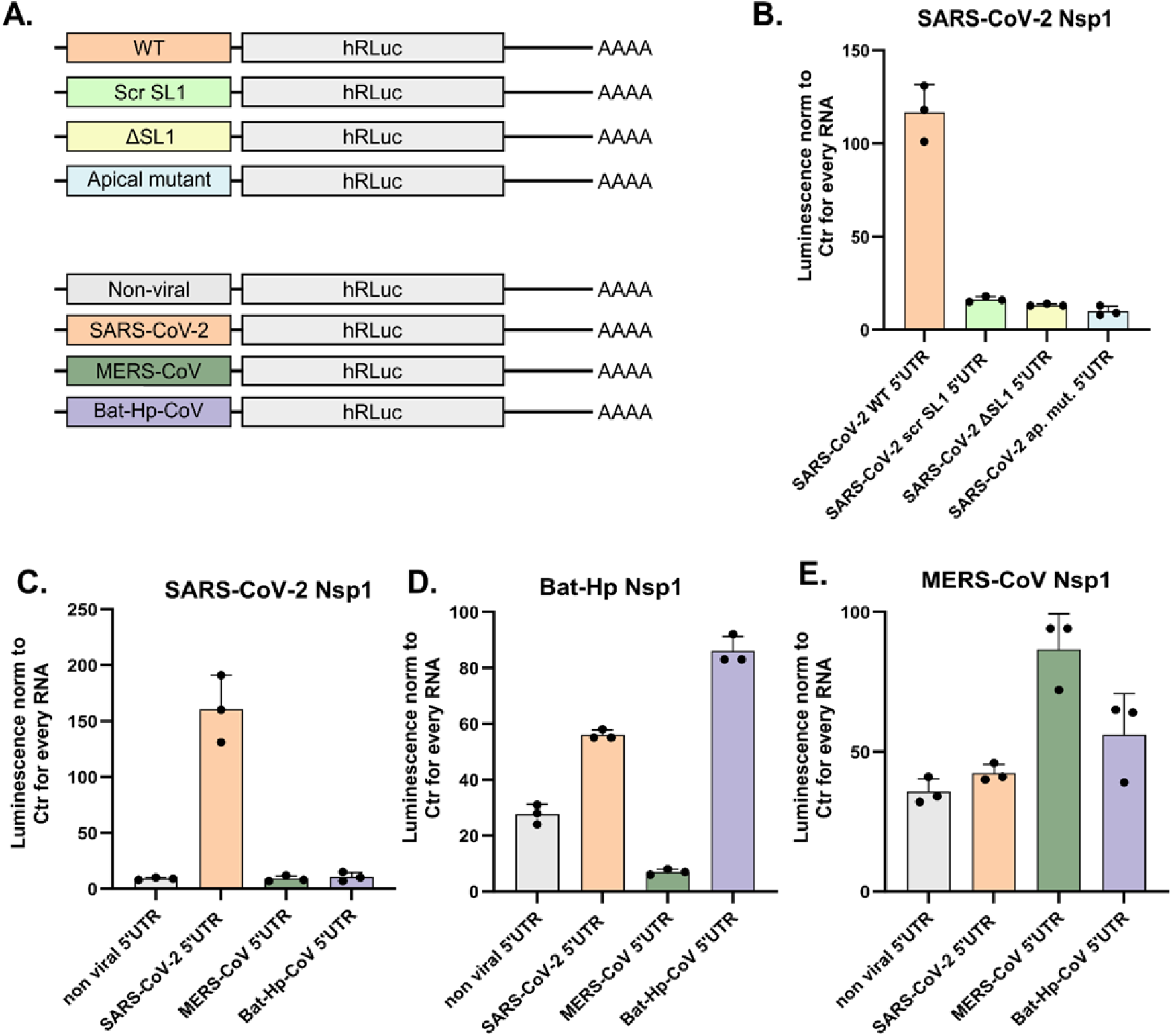
Nsp1 portrays species-specific protection from translation inhibition. **(A)** Schematic representation of Luciferase (RLuc) reporter mRNAs harboring WT and mutant sequences of the full-length SARS-CoV-2 5΄UTR (upper part) and the full-length SARS-CoV-2, MERS-CoV and Bat-Hp-CoV 5΄UTR (lower part). **(B)** Relative RLuc activity measurements of *in vitro* translation reactions supplemented with 0.3 µM of purified SARS-CoV-2 Nsp1 WT, against reporter mRNAs with WT or mutated of SARS-CoV-2 5΄UTRs, each normalized to a reaction using the same RNA and the same concentration of a control protein. **(C)** Relative RLuc activity measurements of *in vitro* translation reactions with 0.3 µM of purified SARS-CoV-2 Nsp1 WT, against reporter mRNAs with WT SARS-CoV-2, MERS-CoV or Bat-Hp CoV 5΄UTRs, each normalized to a reaction using the same RNA and the same concentration of a control protein **(D)** Relative RLuc activity measurements of *in vitro* translation reactions with 0.3 µM of purified Bat-HP-CoV-Nsp1 WT, against reporter mRNAs with WT SARS-CoV-2, MERS-CoV or Bat-Hp CoV 5΄UTRs each normalized to a reaction using the same RNA and the same concentration of a control protein. **(E)** Relative RLuc activity measurements of *in vitro* translation reactions with 0.3 µM of purified MERS-CoV Nsp1 WT, against reporter mRNAs with WT SARS-CoV-2, MERS-CoV or Bat-Hp CoV 5΄UTRs, each normalized to a reaction using the same RNA and the same concentration of a control protein. Data in panels B-E is presented as mean values of three biological replicates (sets of translation reactions) averaged after three measurements.

We compared the activity of 0.3 µM of WT SARS-CoV-2 Nsp1 using equimolar amounts of the different reporter mRNAs in *in vitro* translation reactions. As shown in Fig. 7B, Nsp1 boosts the translation of the RLuc reporter with the WT viral 5΄UTR. However, all three mutant 5’UTRs lead to abolishing the protective effect of the viral 5’UTR against SARS-CoV-2 Nsp1, recapitulating previous observations^10,23,34,35,37^.

Since our *in vitro* system responds to the protective features of the viral 5΄UTR, we set out to assess whether 5΄UTRs originating from genomic RNAs of other Betacoronaviruses have a protective function against SARS-CoV-2 Nsp1. To address this question, we incubated WT Nsp1 with reporter mRNAs harbouring viral 5΄UTRs from SARS-CoV-2, MERS-CoV, and the Bat Hp-CoV (Fig. 7A, lower panel and Fig. 7C). We observed that only the SARS-CoV-2 5’UTR can protect mRNAs from the translation inhibitory effect of the SARS-CoV-2 Nsp1. When we incubated SARS-CoV-2 Nsp1 with reporter mRNAs harbouring the other two 5΄UTRs, we observed no protection (Fig. 7C).

When we performed similar experiments using Bat-Hp or MERS-CoV Nsp1 against reporter mRNAs with 5΄UTRs originating from the other two viral counterparts respectively, we observed that each viral Nsp1 protects mainly reporter mRNAs harbouring the corresponding 5΄UTR, and only partially or not at all reporters with viral 5΄UTRs from different coronaviruses. (Fig. 7E). This result suggests that Nsp1 is likely to have co-evolved with its genomic 5΄UTR to provide protection – or even boost, in the case of SARS-CoV-2 – viral translation in the presence of Nsp1.

## Discussion

To determine whether β-CoV Nsp1-mediated mRNA degradation requires active translation and whether degradation is a secondary effect of translation inhibition, we used a cell-free translation system to decouple the two activities. This study unveils that SARS-CoV-2 Nsp1-dependent mRNA degradation does not depend on active translation and that the two activities can be decoupled.

Different lines of evidence agree that Nsp1 binding to a ribosome is required to stimulate host mRNA degradation ^1^. Intrigued by the elusive mechanism of ribosome-dependent mRNA degradation *in vivo*, we explored whether Nsp1 binding to the 40S ribosomal subunit stimulates Ribosome Quality Control (RQC) mechanisms that could explain the global degradation of host mRNAs. Expression of Nsp1 in HEK-293 cells did not lead to the accumulation of EDF1, a characteristic marker of RQC activation in polysome fractions, weakening the scenario that Nsp1 leads to global ribosome collisions. This observation agrees with biochemical evidence that Nsp1 reduces ribosome stalling of Amyloid Precursor Protein C-Terminal Fragment (APP.C99) ^38^. The finding that Nsp1-mediated degradation does not rely on ribosome collisions prompted us to investigate whether Nsp1-mediated mRNA degradation requires active translation or if binding of Nsp1 to the 40S subunit suffices to degrade host cell mRNAs. Nsp1 translation inhibition and host mRNA degradation are two well-reported activities in living cells, but it is not clear if one depends on the other and whether mRNA degradation requires actively translating ribosomes to ensue ^29^. To decouple the two activities, we used our custom-made cell-free translation system, which is produced by dual centrifugation and allows the reproducible production of ample amounts of human translation-competent lysates. Moreover, applying cell-free translation using lysates of human origin avoids the artefacts that often accompany the widely used *in vitro* translation system from Rabbit Reticulocyte Lysates ^39,40^. Similarly to previous studies, we observed that Nsp1 binding to the ribosome is a prerequisite for stimulating mRNA degradation ^7,16^ because an Nsp1 ribosome-binding mutant (KH-AA) cannot promote degradation. Using different probes, we showed that degradation occurs towards the 5΄end of the transcripts and that the presence of a viral 5΄UTR renders the transcripts immune to Nsp1-dependent degradation, in agreement with previous observations ^17,21^. These findings support that the *in vitro* translation system can recapitulate key elements of Nsp1-dependent mRNA degradation. Using an Nsp1 mutant that primarily affects mRNA degradation (RK-AA) ^11^, we further showed that translation inhibition by Nsp1 does not necessarily lead to mRNA degradation.

Because our cell-free lysates recapitulated previous findings from living cell experiments, we further explored if Nsp1 degradation depends on active translation. We found that Nsp1 stimulates mRNA degradation independently of ribosomal scanning, shown by Nsp1-induced reporter mRNA degradation in the presence of the scanning inhibitor rocaglamide. Furthermore, uncapped mRNA was also degraded in the presence of Nsp1, further supporting that Nsp1-mediated degradation does not depend on canonical initiation. Experiments with the elongation inhibitor puromycin did not affect the degradation of reporter mRNAs in the presence of Nsp1, leading to the conclusion that active translation is not a prerequisite for Nsp1-mediated mRNA degradation and its interaction with the ribosome is sufficient to initiate mRNA degradation. In retrospective, the finding that Nsp1 does not depend on active translation of the targeted mRNA further supports the finding that degradation does not rely on ribosome collisions. Interestingly, while we found that MERS-CoV Nsp1 inhibits host cell translation, explained by its ribosome-binding capacity similar to SARS-CoV-2 Nsp1, it did not lead to mRNA degradation. This finding is significant as it suggests potential mechanistic differences in how MERS-CoV Nsp1 mediates host cell gene expression inhibition compared to SARS-CoV-2.

Overall, our results highlight that our Nsp1-mediated translation inhibition and mRNA degradation are separable processes with distinct mechanisms across different coronaviruses that can be decoupled and studied in cell-free experiments. Experiments from SARS-CoV Nsp1 showed that cleavage can occur either following the formation of ribosomal 48S preinitiation complexes on mRNAs with short (11nt-long) 5’UTRs, or in the presence of GMP-PNP, the nucleotide analogue, which locks the 48S on the initiation codon ^18,19^. This evidence implies that also in SARS-CoV, Nsp1-mediated degradation occurs independently of ribosome collision and does not require active translation.

We tested the previously proposed Nsp1 inhibitors ametantrone and montelukast sodium hydrate. Still, neither compound influenced the capacity of Nsp1 to inhibit translation, suggesting that the quest for potential Nsp1 inhibitors is still ongoing. In accordance to our findings, previous work showed that Montelukast did not lead to Nsp1 inhibition in cell culture experiments ^16^ nor *in vitro* RNA degradation assays ^21^.

Last, we observe, especially for SARS-CoV-2, that reporter RNAs harbouring a 5΄UTR from the same virus are immune to Nsp1 activities in agreement with a previous study suggesting that the RNA-protein interplay that fine-tunes Nsp1 function is evident in SARS-CoV-2 ^5^. Our comparative analysis indicates a possible co-evolution of SL1 sequences with Nsp1 to evade translation inhibition through a mechanism that implicates the N-terminal domain of Nsp1 ^9^.

It remains unclear which cellular factors are essential for Nsp1-mediated endonucleolytic cleavage of non-viral mRNAs during infection. In a reconstituted *in vitro* system, Nsp1, 40S subunits, and eIF3g alone are sufficient to induce mRNA cleavage ^21^. The host cell exonuclease Xrn1 triggers 5′–3′ mRNA degradation following the expression of SARS-CoV Nsp1 ^22^ and interactome studies on SARS-CoV-2 Nsp1 suggest that it associates with various cellular endo- and exonucleases ^23^.Our results highlight that the two major activities of Nsp1, host mRNA translation inhibition and degradation, can be decoupled, but both require ribosome binding to stimulate host gene expression inhibition. For this reason, developing Nsp1 inhibitors should focus on the interaction between Nsp1 and the 40S subunit.

### Limitations of the study

While this work provides valuable insights into the mechanistic differences of different Nsp1 proteins from SARS-CoV-2 and MERS-CoV, our study relies primarily on a cell-free translation system, which, although highly informative and controllable, may not fully recapitulate the complexity of intracellular environments. For instance, the interaction between Nsp1 and host factors and the impact of cellular compartmentalization could differ in living cells. While we observe that SARS-CoV-2 Nsp1 induces mRNA degradation independently of ribosome collisions, our data does not exclude the possibility that ribosome collisions may occur to a low extent in the presence of Nsp1 nor that they may minimally contribute to host mRNA degradation. Moreover, although our findings on viral mRNA protection by Nsp1 suggest co-evolutionary adaptations, we did not test the functional implications of this protection in cases of co-infection or across a broader spectrum of coronaviruses.

## Acknowledgements

This work was supported by EDK grants from the Swiss National Science Foundation (SNSF CRSK-3_220624), the Multidisciplinary Center of Infectious Diseases from the University of Bern (MCID), the Fondation Claude et Giuliana, the Forschungsstiftung of the University of Bern and the National Center of Excellence in Research (NCCR) on RNA and Disease funded by the SNSF (51NF40-205601). We thank Erik Basha and Sarah Jordi for preliminary results and Jana Ziegelmüller for critically reading the manuscript. Figure 2A and the graphical abstract were made using Biorender.

## Author contributions Declaration of Interests

E.B., D.A., J.L, A.L and N.S. conducted the experiments; E.K. designed the experiments and wrote the initial manuscript, all authors read and edited the manuscript.

## Supplemental information

Document S1. Figures S1–S8

## Supplementary figures

**Fig. S1.**
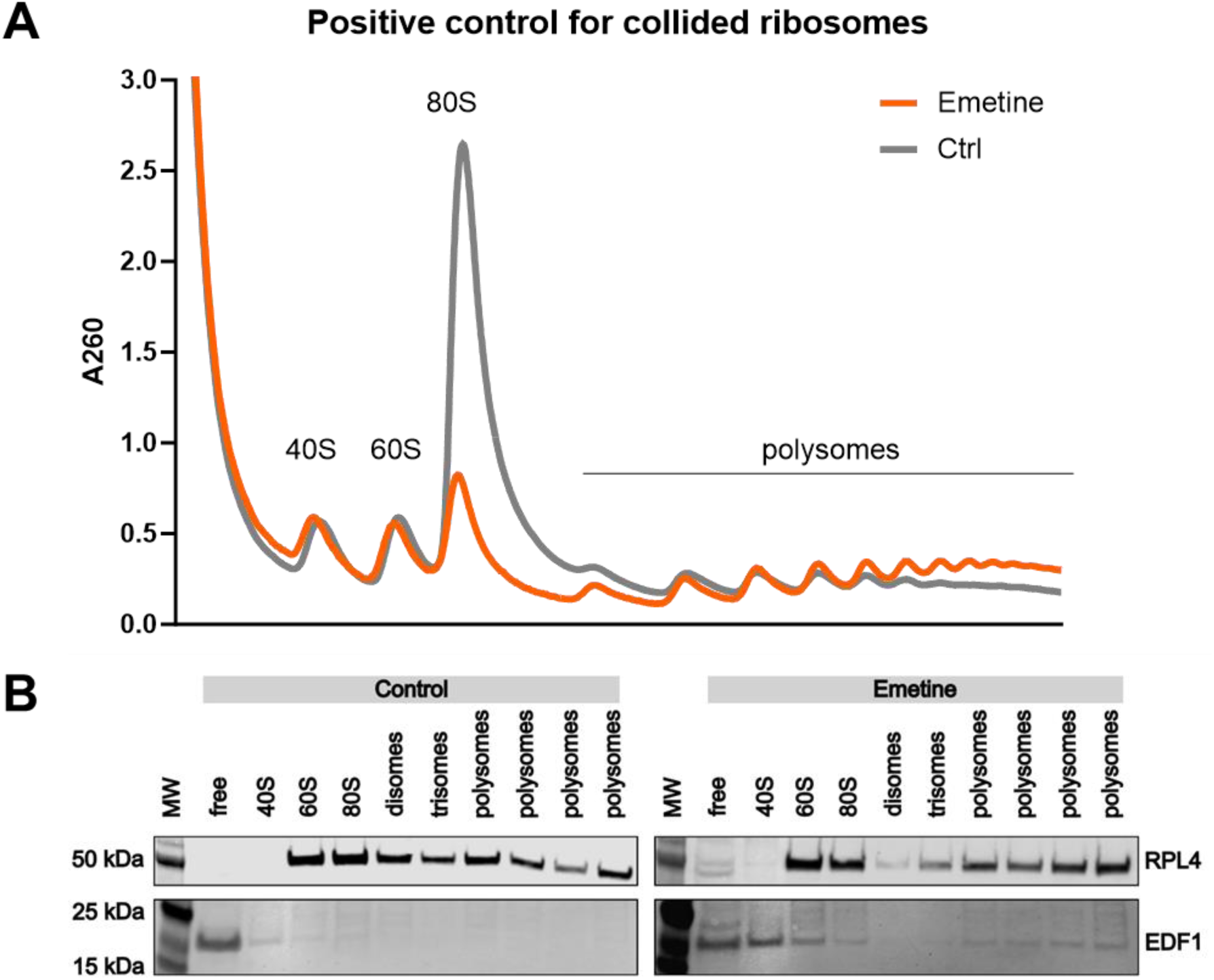
related to Fig. 1. Positive control for collided ribosomes. **(A)** Equal amounts of untreteated (control) and emetine-treated cell lysates from HEK293T cells were fractionated on a 15-50% sucrose gradient by ultracentrifugation. **(B)** From the collected polysome fractions, total proteins were extracted with acetone and TCA. Detected proteins are labelled on the right side of the blots.

**Fig. S2.**
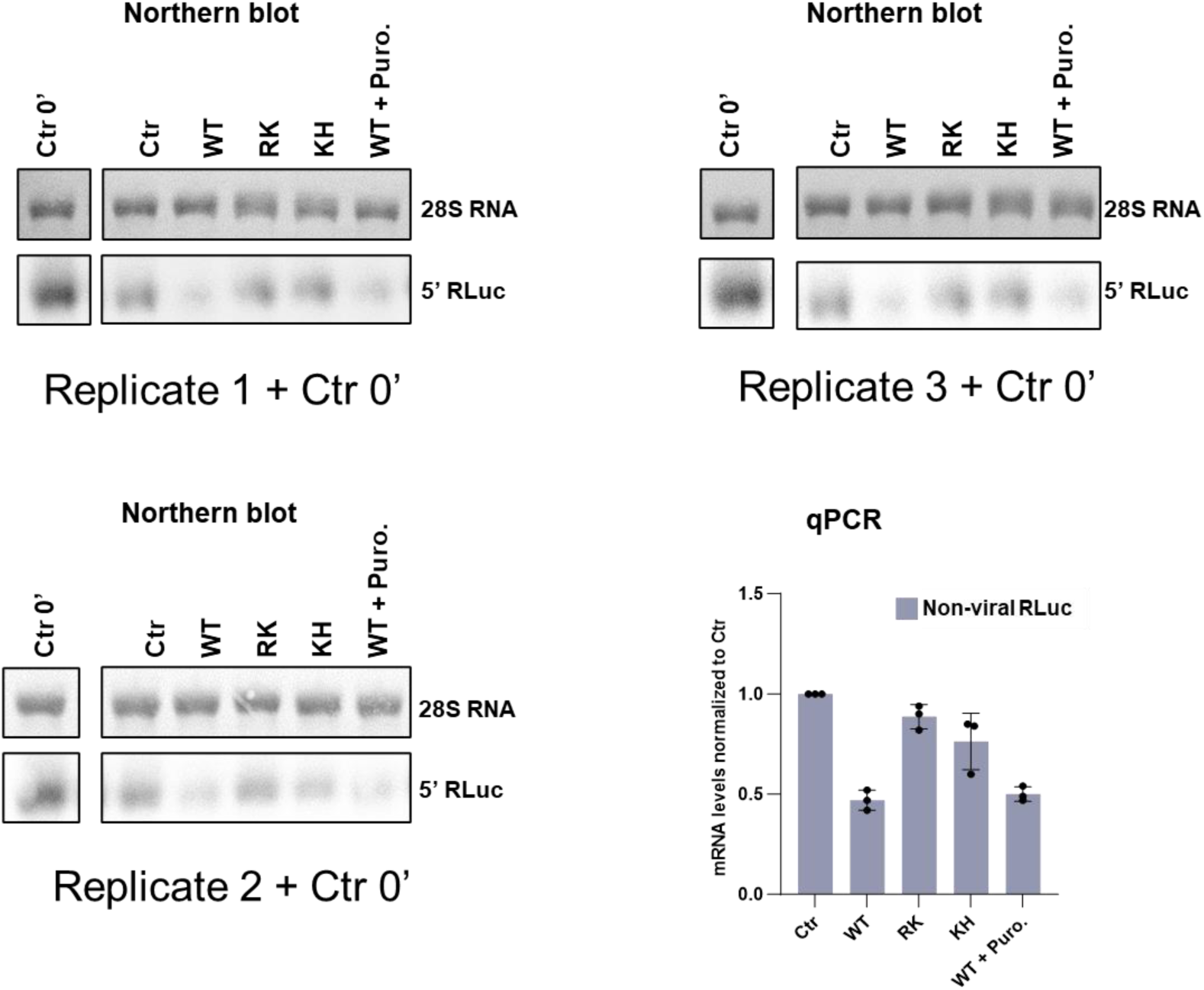
related to Fig. 2C. Three replicates of Northern blot referring to panel 2C including the 0΄control aside. 500 ng of isolated RNA from the *in vitro* translation reactions was subjected to Northern blot analysis using a 32P-labelled probe for the reporter RLuc RNA. The methylene blue-stained pattern of the 28S rRNA is shown as an internal standard for RNA loadings for each lane. Data in the quantification plot are presented as mean values of three biological replicates (sets of translation reactions) averaged after three densitometry measurements. qPCR - Relative mRNA levels normalized to the Ctr reaction determined by RT-qPCR and normalized to the average of the relative levels of the normalizer RNA.

**Fig. S3.**
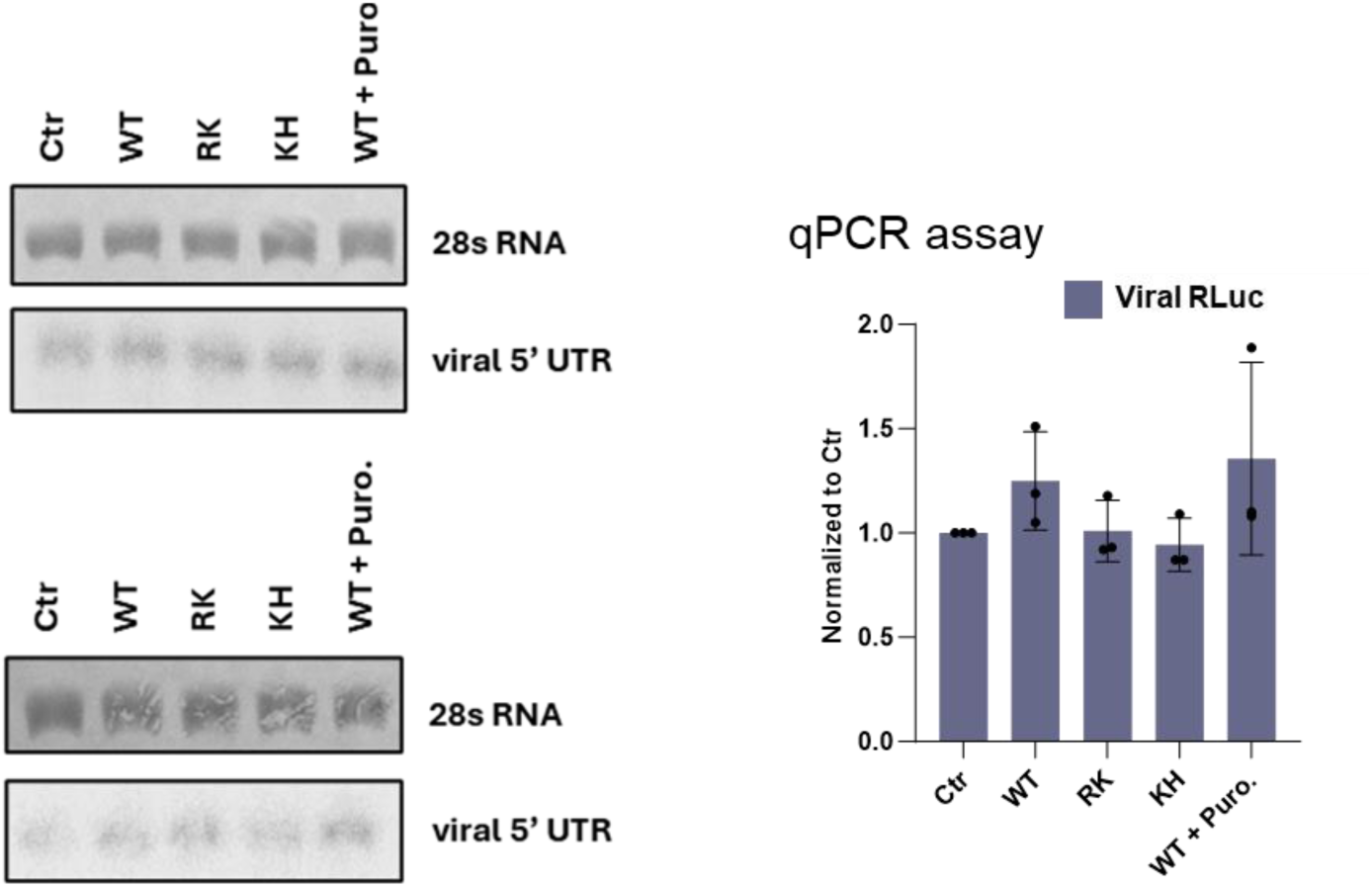
related to Fig. 2F. Two additional replicates of Northern blot referring to panel 2F. 500 ng of isolated RNA from the *in vitro* translation reactions was subjected to Northern blot analysis using a 32P-labelled probe for the reporter RLuc RNA. The methylene blue-stained pattern of the 28S rRNA is shown as an internal standard for RNA loadings for each lane. Data in the quantification plot are presented as mean values of three biological replicates (sets of translation reactions) averaged after three densitometry measurements. qPCR - Relative mRNA levels normalized to the Ctr reaction determined by RT-qPCR and normalized to the average of the relative levels of the normalizer RNA.

**Fig. S4.**
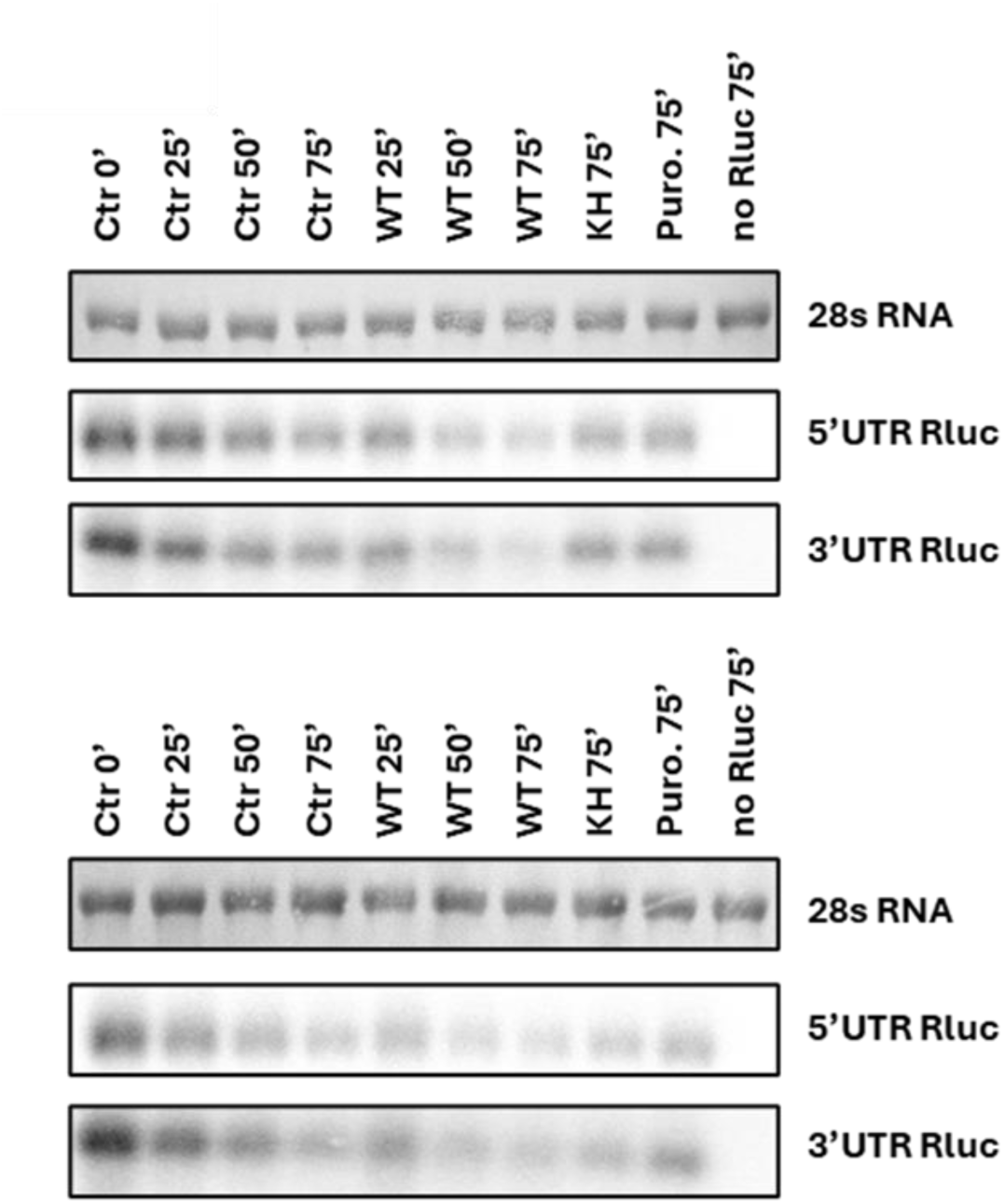
related to Fig. 3C. Two additional replicates of Northern blot referring to Fig. 3C. 500 ng of isolated RNA from the *in vitro* translation reactions was subjected to Northern blot analysis using a 32P-labelled probe for the reporter RLuc RNA. The methylene blue-stained pattern of the 28S rRNA is shown as an internal standard for RNA loadings for each lane.

**Fig. S5.**
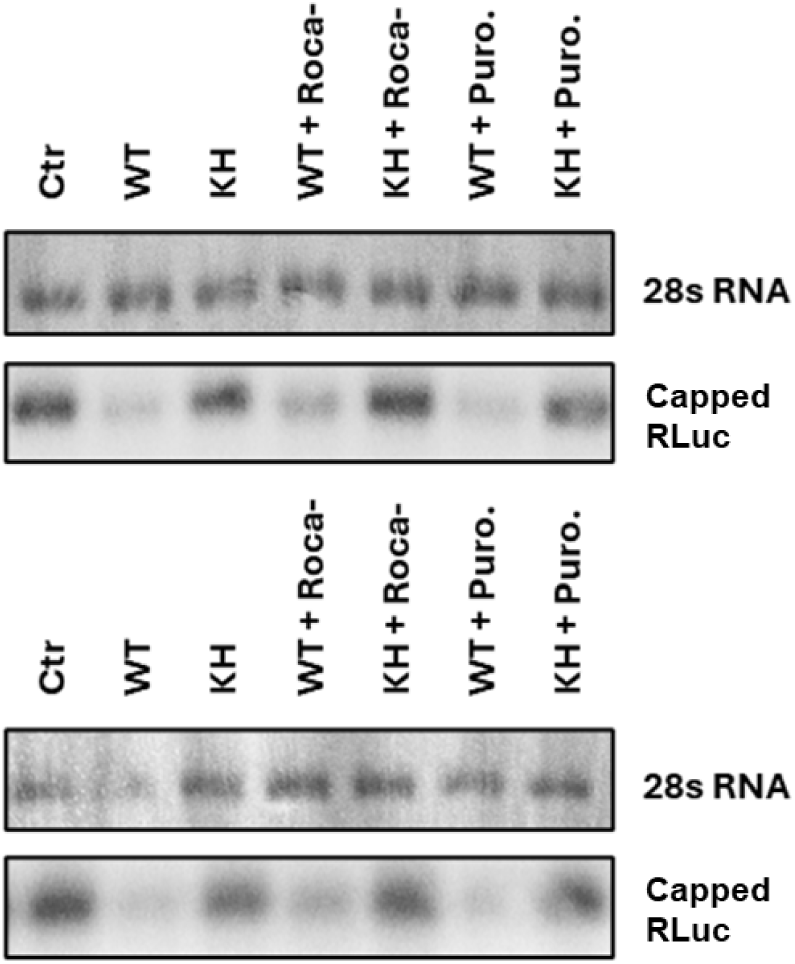
related to Fig. 4B. Two additional replicates of Northern blot referring to Fig. 4B. 500 ng of isolated RNA from the *in vitro* translation reactions was subjected to Northern blot analysis using a 32P-labelled probe for the reporter RLuc RNA. The methylene blue-stained pattern of the 28S rRNA is shown as an internal standard for RNA loadings for each lane.

**Fig. S6.**
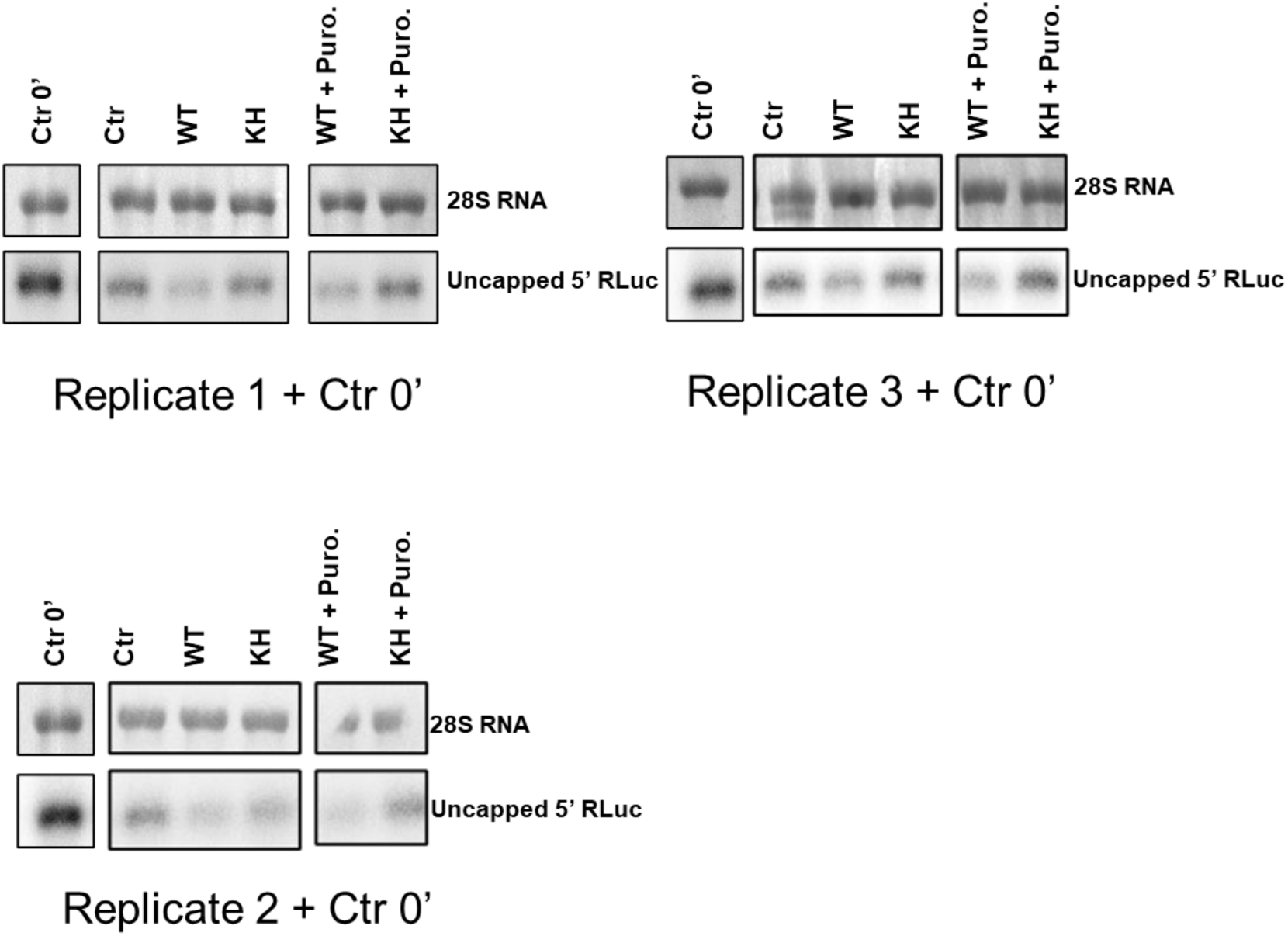
related to Fig. 4D. Three replicates of Northern blot referring to Fig. 4D including the 0΄control aside. 500 ng of isolated RNA from the *in vitro* translation reactions was subjected to Northern blot analysis using a 32P-labelled probe for the reporter RLuc RNA. The methylene blue-stained pattern of the 28S rRNA is shown as an internal standard for RNA loadings for each lane.

**Fig. S7.**
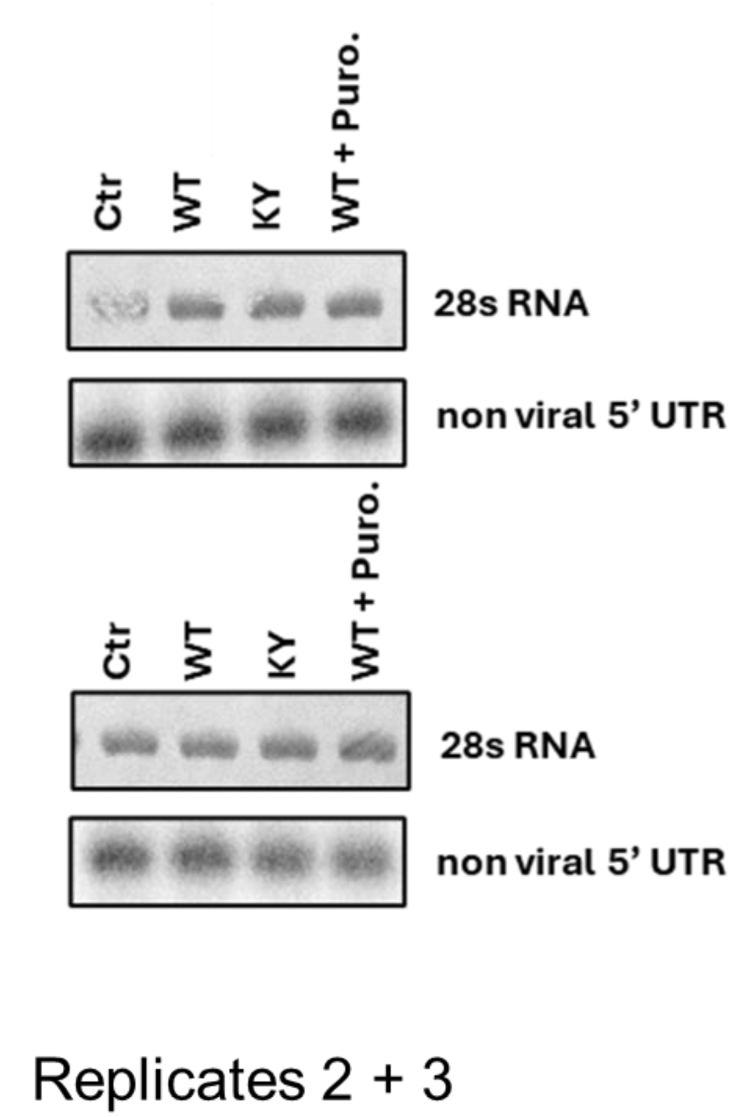
related to Fig. 5B. Two additional replicates of Northern blot referring to Fig. 5B including the 0΄control aside. 500 ng of isolated RNA from the *in vitro* translation reactions was subjected to Northern blot analysis using a 32P-labelled probe for the reporter RLuc RNA. The methylene blue-stained pattern of the 28S rRNA is shown as an internal standard for RNA loadings for each lane.

**Fig. S8.**
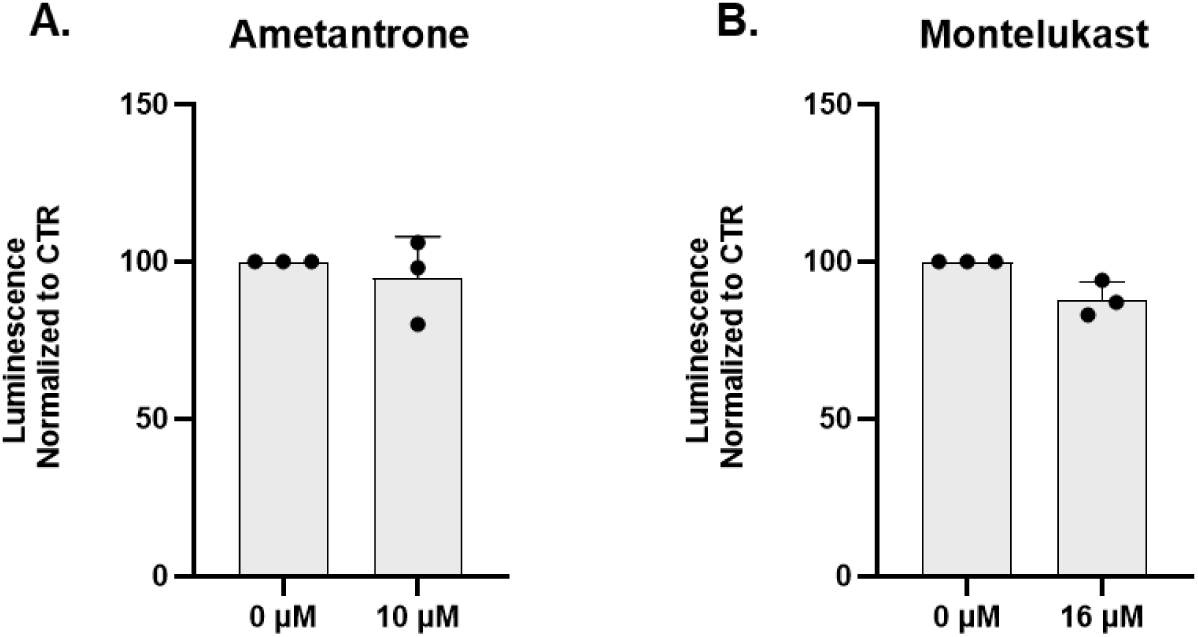
related to Fig. 6. Relative *RLuc* luminescence measurements of *in vitro* translation reactions with maximum concentrations used of Ametantrone **(A)** or Montelukast **(B)** in the without SARS-CoV-2 Nsp1 WT, normalized to the reaction in the presence of no drug. Data are presented as mean values of three biological replicates (sets of translation reactions) averaged after three measurements. The positive control reaction for a translation inhibition contained 1 mg/mL puromycin.

